# Cell-type specific circadian bioluminescence rhythms in *Dbp* reporter mice

**DOI:** 10.1101/2021.04.04.438413

**Authors:** Ciearra B. Smith, Vincent van der Vinne, Eleanor McCartney, Adam C. Stowie, Tanya L. Leise, Blanca Martin-Burgos, Penny C. Molyneux, Lauren A. Garbutt, Michael H. Brodsky, Alec J. Davidson, Mary E. Harrington, Robert Dallmann, David R. Weaver

**Affiliations:** Department of Neurobiology, University of Massachusetts Chan Medical School, Worcester MA; Graduate Program in Neuroscience, University of Massachusetts Chan Medical School, Worcester MA; Department of Biology, Williams College, Williamstown, MA; Neuroscience Program, Smith College, Northampton MA; Neuroscience Institute, Morehouse School of Medicine, Atlanta GA; Department of Mathematics and Statistics, Amherst College, Amherst MA; Division of Biomedical Sciences, Warwick Medical School, University of Warwick, Coventry, UK; Department of Molecular, Cell and Cancer Biology, University of Massachusetts Chan Medical School, Worcester MA; NeuroNexus Institute, University of Massachusetts Chan Medical School, Worcester MA

**Keywords:** Circadian Rhythms, Bioluminescence, Luciferase, Misalignment, *Dbp*, Albumin D-element binding protein, *In Vivo* Imaging System (IVIS), LumiCycle *In Vivo*, Reporter Mouse, Peripheral Oscillators

## Abstract

Circadian rhythms are endogenously generated physiological and molecular rhythms with a cycle length of about 24 h. Bioluminescent reporters have been exceptionally useful for studying circadian rhythms in numerous species. Here, we report development of a reporter mouse generated by modification of a widely expressed and highly rhythmic gene encoding D-site albumin promoter binding protein (*Dbp*). In this line of mice, firefly luciferase is expressed from the *Dbp* locus in a *Cre*-recombinase-dependent manner, allowing assessment of bioluminescence rhythms in specific cellular populations. A mouse line in which luciferase expression was *Cre*-independent was also generated. The *Dbp* reporter alleles do not alter *Dbp* gene expression rhythms in liver or circadian locomotor activity rhythms. *In vivo* and *ex vivo* studies show the utility of the reporter alleles for monitoring rhythmicity. Our studies reveal cell-type specific characteristics of rhythms among neuronal populations within the suprachiasmatic nuclei *ex vivo*. *In vivo* studies show *Dbp*-driven bioluminescence rhythms in the liver of *Albumin-Cre;Dbp^KI/+^* “liver reporter” mice. After a shift of the lighting schedule, locomotor activity achieved the proper phase relationship with the new lighting cycle more rapidly than hepatic bioluminescence did. As previously shown, restricting food access to the daytime altered the phase of hepatic rhythmicity. Our model allowed assessment of the rate of recovery from misalignment once animals were provided with food *ad libitum*. These studies confirm the previously demonstrated circadian misalignment following environmental perturbations and reveal the utility of this model for minimally invasive, longitudinal monitoring of rhythmicity from specific mouse tissues.

## Introduction

Circadian rhythms are endogenous rhythms with a cycle length of ∼24 hours. The mammalian circadian system is hierarchical, with the hypothalamic suprachiasmatic nuclei (SCN) serving as the pacemaker (Mohawk et al., 2012; Herzog et al., 2017). The SCN are synchronized by environmental cues, of which the light-dark cycle is the most influential. The SCN are not unique in their capacity for rhythmicity, however. The transcriptional-translational feedback loop regulating molecular oscillations in the SCN is also present in individual cells throughout the body (Mohawk et al., 2012). SCN-driven neural, behavioral and hormonal rhythms synchronize these cell-autonomous oscillators, leading to rhythmicity with predictable phase relationships among tissues, genes and physiological processes (Mohawk et al., 2012; Patke et al., 2020; Zhang et al., 2014). Repeated disruption of this internal temporal order by inappropriately timed light exposure or food intake leads to adverse health consequences in shift-working humans and in animal models (Evans & Davidson, 2013; Patke et al., 2020). Progress in identifying the mechanisms by which chronic circadian disruption leads to adverse health consequences will require long-term monitoring of central and peripheral rhythms (Roenneberg & Merrow, 2016).

Rhythmically expressed reporter genes have been extremely important for demonstrating cell-autonomous circadian clocks and monitoring rhythmicity in several organisms, including plants (Millar et al., 1992), *Neurospora* (Morgan et al., 2003), cyanobacteria (Kondo et al., 1993), *Drosophila* (Brandes et al., 1996), zebrafish (Weger et al., 2013), cultured cells (Nagoshi et al., 2004; Hirota et al., 2010; Welsh et al., 2004; Zhang et al., 2009), rodent tissue explants (Abe et al., 2002; Maywood et al., 2013; Yamazaki et al., 2000; Yoo et al., 2004; Yoo et al., 2005), and rodent tissues *in vivo* (Saini et al., 2013; Tahara et al., 2012). Circadian reporter genes have been instrumental in screens to identify clock genes and modifiers in many of these systems (Cesbron et al., 2013; Chen et al., 2012; Hirota et al., 2010; Kondo et al., 1993; Millar et al., 1995; Muñoz-Guzmán et al., 2021; Stanewsky et al., 1998; Zhang et al., 2009). Circadian reporters have also been used to assess rhythmicity in peripheral tissues and the impact of alterations in experimental or environmental conditions (food availability, lighting cycles, glucocorticoid treatment) on peripheral oscillators, conducted by measuring bioluminescence rhythms in tissue explants monitored *ex vivo* (Davidson et al., 2008; Davidson et al., 2009; Nakamura et al., 2005; Pezuk et al., 2012; Sellix et al., 2012; Stokkan et al., 2001; Yamanaka et al., 2008; Yamazaki et al., 2000). These studies complement work done by assessing population rhythms in gene expression in tissue samples indicating altered rhythm amplitude and phase, and altered phase relationships in and between SCN and peripheral oscillators following resetting (Balsalobre et al., 2000; Damiola et al., 2000; Destici et al., 2013;Nagano et al., 2003; Reddy et al., 2002; Yamaguchi et al., 2013; for review see Nicholls et al., 2019). Several groups have developed methods for *in vivo* assessment of reporter gene activity from brain regions, including the SCN, using implanted optical fibers and freely moving (but tethered) rodents (Hamada et al., 2016; Mei et al., 2018; Nakamura et al., 2008; Ono et al., 2015; Yamaguchi et al., 2001; Yamaguchi et al., 2016). Other studies have localized the source of bioluminescence from widely expressed reporter genes in specific peripheral tissues based on photomultiplier tube placement on the body surface (Hamada et al., 2016; Sawai et al., 2019). Peripheral organ reporter gene activity has been assessed by *in vivo* imaging in anesthetized mice (Saini et al., 2013; Tahara et al., 2012) and more recently in ambulatory mice (Martin-Burgos et al., 2020; Saini et al., 2013; Sinturel et al., 2021). In some cases, viral vectors that afford anatomical specificity (through their site of injection, tropism and/or by their design) have been used to direct reporter expression to specific tissues (Mei et al., 2018; Saini et al., 2013; Sinturel et al., 2021). All of these approaches are hampered by the need to develop specific reagents or approaches for each tissue being examined, and many of these approaches are invasive. In view of the large number of mouse lines with tissue-specific expression of *Cre* recombinase, the field would benefit considerably from a binary (*Cre*-lox) reporter system in which bioluminescence from a rhythmically expressed gene can be switched on in tissues expressing *Cre* recombinase, simply by crossing mice of the appropriate genotypes together.

Here, we report a new transgenic mouse line in which firefly luciferase is expressed from the mouse *Dbp* locus in a *Cre*-recombinase-dependent manner. *Dbp* is widely and rhythmically expressed (Fonjallaz et al., 1996; Punia et al., 2012; Zhang et al., 2014), allowing detection of circadian bioluminescence rhythms in numerous tissues, *in vivo* and *ex vivo*. *Cre*-dependent bioluminescence rhythms were recorded *ex vivo* from specific SCN neuronal populations. Furthermore, we observed transient misalignment between behavioral and hepatic bioluminescence rhythms in freely moving mice subjected to a shift of the light-dark cycle or following restricted food access.

While this work was being prepared for publication, Shan et al. (2020) reported development of a Color-Switch *Per2* reporter mouse. In this reporter, *Cre* recombinase expression changes the reporter fused to mPER2 from red to green luciferase.

## Materials and Methods

### Animals and Housing Conditions

All animal procedures were reviewed and approved by the Institutional Animal Care and Use Committees of the University of Massachusetts Chan Medical School, Morehouse School of Medicine, the University of Warwick, and/or Smith College.

Unless otherwise noted, animals were maintained in a 12h light: 12h dark (LD) lighting cycle with access to food (Prolab Isopro RMH3000; LabDiet) and water available *ad libitum*. Zeitgeber Time (ZT) refers to time relative to the lighting cycle. ZT 0-12h is the light phase and ZT 12-24h is the dark phase.

*Cre* recombinase-expressing lines were crossed to mice bearing the conditional (*Dbp^KI^*) reporter allele to generate mice expressing luciferase in specific cells or tissues. *Albumin-Cre* (B6.Cg-*Speer6-ps1^Tg(Alb-Cre)21Mgn^*/J; JAX stock number 003574), *Ksp1.3-Cre* (B6.Cg-Tg{Cdh16-cre}91Igr/J, JAX 012237), *AVP*-IRES2-*Cre* (B6.Cg-*Avp^tm1.1(Cre)Hze^*/J; JAX 023530), and *NMS*-*Cre* mice (Tg(Nms-iCre)^20Ywa^, JAX 027205) were obtained from the Jackson Labs (Bar Harbor, ME). These lines direct *Cre* recombinase expression to hepatocytes (Postic et al., 1999), renal tubules and genito-urinary epithelia (Shao et al., 2002), neurons expressing arginine vasopressin (AVP; Harris et al., 2014), and neurons expressing Neuromedin S (NMS; Lee et al., 2015), respectively. A *Prrx1-Cre* female (B6.Cg-Tg(Prrx1-Cre^1Cjt^/J), JAX 005584; Logan et al., 2002) was used for germline deletion of the conditional allele (see below).

Founder *Per2^LucSV/+^* mice with an in-frame fusion of firefly luciferase to PER2 and an SV40 polyadenylation signal (Welsh et al., 2004; Yoo et al., 2017) were generously provided by Dr. Joseph Takahashi, University of Texas Southwestern Medical School, Dallas. All *Per2^LucSV^* reporter mice used for experiments here were heterozygous (e.g., *Per2^LucSV/+^*). For clarity when referring to literature describing the more widely used *PER2::LUCIFERASE* fusion reporter line in which the endogenous *Per2* 3′ UTR is downstream of the luciferase coding sequence (Yoo et al., 2004), we will refer to this line as *Per2^Luciferase^.* Mouse lines were maintained by backcrossing to the C57BL/6J (JAX 000664) background.

We also generated albino reporter mice by backcrossing to albino C57BL/6J mice with a mutation in tyrosinase (*tyr/tyr*; B6(Cg)-*Tyr^c-2J^*/J, JAX stock number 00058). Tyrosinase, like *Dbp*, is located on mouse chromosome 7. Crossing these lines eventually generated a recombinant (*Dbp^KI/+^; tyr/tyr*) in which both mutant alleles were on the same chromatid. Subsequent crossing to albino mice expressing *Cre* recombinase allowed production of albino reporter mice. Albino *Dbp^KI/+^* mice on the B6(Cg)-*Tyr^c-2^*J/J background are being deposited in the Jackson Labs repository (Bar Harbor, ME) as stock number 036997.

Note, caution is needed with the *Ksp1.3-Cre* line reported here, as it has a high frequency of germline recombination (excision of the floxed region of the conditional allele in the germline, leading to non-conditional luciferase expression) when *Ksp1.3-Cre* is present in the same parent as *Dbp^KI/+^*. Recombination also frequently occurs when *Ksp1.3-Cre* females are crossed with *Dbp^KI^* males. When using the *Cre/lox* system, genotying strategies should be designed to detect all possible alleles. Even when the *Ksp1.3-Cre; Dbp^KI/+^* genotype is generated without germline excision of GFP, the sex difference in *Cre* expression leads to markedly different bioluminescence patterns in males and females (see Results).

### CRISPR/Cas9 targeting the *Dbp* locus

The mutant allele was generated by CRISPR/Cas9 mediated engineering of the *Dbp* locus. The targeting construct (**Figure 1**) consisted of a 5′ homology arm terminating just 5′ of the *Dbp* stop codon followed by in-frame sequences encoding a T2A linker (to separate DBP protein from the reporter polypeptides; Kim et al., 2011), loxP, GFP with the bovine growth hormone polyadenylation signal, loxP, and *Luc2* followed by the 3′-UTR of *Dbp* (3′ homology arm). In the presence of CRE recombinase, two loxP sites oriented in the same direction will recombine, leading to deletion of the sequence between them (GFP in this case).

**Figure 1.**
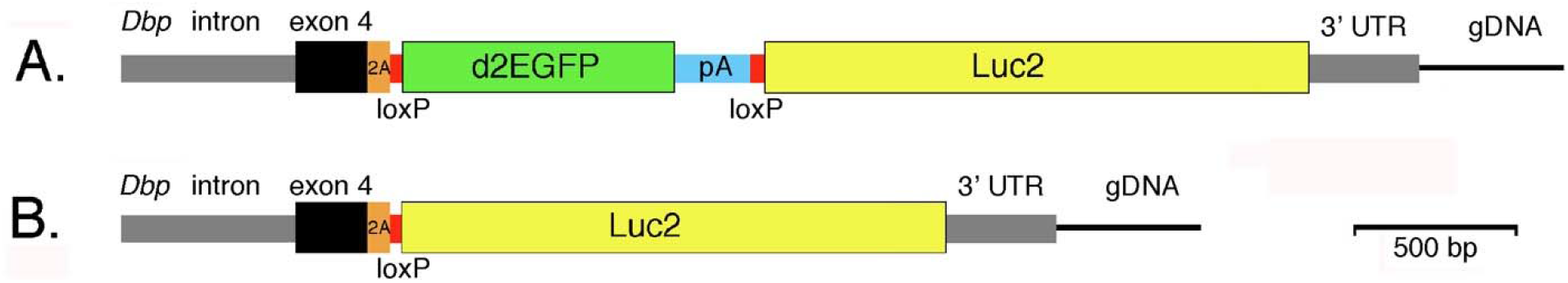
Generation of a bifunctional reporter from the mouse *Dbp* locus. **A.** The mouse *Dbp* locus was modified by CRISPR-mediated insertion of the donor construct shown. The construct contained homology arms from the *Dbp* locus (gray and black) and inserted the reporter sequences with a T2A-encoding sequence (orange) between DBP and the reporter. Destabilized EGFP (d2EGFP) with a bovine growth hormone polyadenylation site (PA) was flanked by *loxP* sites (red). Downstream of *GFP* is a luciferase (*Luc2*) reporter gene. Without recombination *Dbp* and *GFP* are expressed as a single transcript from the conditional (*Dbp^KI^* allele). **B.** With *Cre*-mediated recombination, GFP-encoding sequences are excised and *Dbp* and *luciferase* are expressed as a single transcript. The T2A sequence generates separate proteins from these bifunctional transcripts. *Cre*-mediated germline recombination led to mice expressing luciferase non-conditionally from the *Dbp^Lu^*^c^ allele.

In the successful set of microinjections, 34 blastocysts were injected with 40 ng/μl guide RNA MmDBPki_gR49f, 50 ng/μl *Cas9* mRNA (synthesized from a Cas9 PCR product using mMessage mMachine T7 Ultra Kit from Life Technologies) and 20 ng/μl CAS9 protein (IDT). Two putative founders were identified using a primer pair internal to the construct (primer pair C; **Table S1**). Additional primer pairs consisting of a primer in flanking DNA (external to the construct) and a primer within the construct were used to determine whether these animals had the desired targeting event (primer pairs F and H, which spanned the 5′ and 3′ ends, respectively). These studies led to identifying one mouse as having the correct insertion and recognizing that the other putative founders had random insertion of the construct rather than homologous recombination into the *Dbp* locus; the mouse with random insertion was not studied further. Genomic DNA from the founder with insertion into the *Dbp* locus was amplified using a primer pair flanking the entire construct. Sequencing the product confirmed the construct was inserted properly, *in vivo*. Primer sets used for verification of the proper insertion of the construct are listed in **Table S1**.

The founder carrying the targeted (knock-in or *Dbp^KI^*) allele and its offspring were backcrossed to C57BL/6J mice (JAX 000664) for three generations before any intercrossing to reduce the chance of a potential off-target mutations becoming established in the reporter line.

To generate mice with germline deletion of GFP (and thus leading to expression of luciferase throughout the body), a male *Dbp^KI/+^* was bred to a *Prrx1-Cre* female, which we had on hand and which, in our experience, produces germline deletion of floxed alleles at high frequency when CRE is introduced from the female. Several mice bearing the newly generated *Dbp^Luc^* allele were identified and backcrossed to C57BL/6J mice, selecting against *Prrx1-Cre*.

### Genotyping

Genotyping was performed by PCR amplification of DNA extracted from ear punches. Amplification products were separated by agarose gel electrophoresis. Genotyping protocols for *Per2^LucSV^* and *Cre* recombinase have been published previously and are listed in Table S1 (van der Vinne et al., 2018; Weaver et al., 2018, respectively). A mixture of four primers (primer set “4A”) capable of detecting all possible *Dbp* allele combinations was used for colony genotyping; the three possible alleles (*Dbp^KI^*, *Dbp^Luc^*, *Dbp^+^*) generate amplicons of 399, 490 and 299 bp, respectively with this primer set. Primer set 4A consists of a common forward primer in exon 4 (5′-TGCTGTGCTTTCACGCTACCAGG-3′) and allele-specific reverse primers in GFP (to detect the *Dbp^KI^* allele; 5′-AGTCGTGCTGCTTCATGTGGTCG-3′), in Luc2 (to detect the *Dbp^Luc^* allele; 5′-TCGTTGTAGATGTCGTTAGCTGG-3′), and in the *Dbp* 3′ UTR (to detect the unmodified *Dbp* allele; 5′-TTCAGGATTGTGTTGATGGAGGC-3′).

### Generation of Digoxigenin (DIG) DNA Probes and Northern Blot Assay

DIG-labeled DNA probes were generated by PCR in reactions containing 28 μM of DIG-labeled UTP. Primer sets are listed in **Table S1**.

Male mice of five genotypes (WT, *Dbp^KI/+^*, *Dbp^KI/KI^*, *Dbp^Luc/+^*, and *Dbp^Luc/Luc^*) were euthanized by Euthasol injection for collection of liver tissue at 4-h intervals (ZT 2, 6, 10, 14, 18, 22). RNA was isolated from the liver tissue by Trizol extraction (Ambion). RNA was quantitated by Nanodrop. Five micrograms per lane were separated by electrophoresis on 1.2% formaldehyde gels. RNA was transferred to nylon membranes and cross-linked by UV exposure. Blots were prehybridized, probed and detected following the manufacturer’s protocol (Roche), bagged and exposed to X-ray film.

Film images of the blots were analyzed by determining the optical density of the *Dbp* and *Actin* bands within each lane and taking the *Dbp/Actin* ratio. The *Dbp/Actin* ratios were converted to percentage of maximum *Dbp/Actin* for each transcript type within each blot. Due to the difference in band location of the three *Dbp* alleles, heterozygous animals contributed a set of values for both the wild-type transcript and the reporter transcript on each blot. Friedman’s one-way analysis of variance and Dunn’s test were used for non-parametric assessment of differences between time-points for each transcript.

### Locomotor Activity Rhythms

Male and female mice of five genotypes (WT, *Dbp^KI/+^*, *Dbp^KI/KI^*, *Dbp^Luc/+^*, and *Dbp^Luc/Luc^*) were transferred to the experimental room and single-housed with a running wheel. Animals had access to food and water *ad libitum*. Running-wheel activity was monitored and analysed using ClockLab collection software (Actimetrics). Mice were entrained to a 12-h light/12-h dark cycle for 18 days, then were placed into constant darkness (dim red light) for 15 days. The free-running period in constant darkness (DD) was determined for each animal on DD days 4-15 by periodogram analysis (ClockLab).

### Bioluminescence Recordings from Tissue Explants

Tissue explants were prepared late in the afternoon from *Per2^LucSV/+^* and *Dbp^Luc/+^* mice housed on a 12-h light/12-h dark lighting cycle. Tissues from the two genotypes were studied together in each run. Mice were deeply anesthetized with Euthasol and decapitated. Tissues were dissected and immediately placed in ice-cold 1X HBSS (Gibco). Pituitary gland was subdivided into 4 sections (∼2mm^3^) with a scalpel and each piece was cultured separately. Lung explants were placed three per dish. Up to three replicate dishes were studied per tissue per animal. Explants were placed on sterile 35-mm Millicell culture plate inserts (Millipore) in a sealed petri dish containing air-buffered bioluminescence medium (Yamazaki and Takahashi, 2005) plus D-luciferin (100 μM) (Gold Biotechnology) and incubated at 32 °C as previously described (van der Vinne et al., 2018). Bioluminescence in each dish was measured for 1 minute every 15 minutes using a Hamamatsu LM-2400 luminometer.

Bioluminescence records were analyzed using Microsoft Excel to determine period and peak time. The first 12-h were discarded to exclude acute responses to explant preparation. Photon counts were smoothed to a 3-h running average and baseline subtracted using a 24-h running average. Circadian period was determined from the average of the period between each peak, trough, upward crossing and downward crossing between 24 and 88 hr of recording for each record. Peak time was calculated as the clock time of the first peak in the background-subtracted data and is expressed relative to ZT of the extrapolated lighting cycle.

### Imaging of Bioluminescence Rhythms *In Vivo*

*In vivo* imaging was performed in the UMass Chan Medical School Small Animal Imaging Core Facility using an *In Vivo* Imaging System (IVIS-100, Caliper, now Perkin Elmer) as previously described (van der Vinne et al., 2018; van der Vinne et al., 2020). *Alb-Cre^+^; Dbp^KI/+^* (liver reporter), *Dbp^Luc/+^*, and *Per2^LucSV/+^* mice were anesthetized with 2% isoflurane (Zoetis Inc.) and skin covering the liver, kidneys and submandibular glands was shaved. Mice were injected with D-luciferin (i.p., 100 μl at 7.7 mM, Gold Biotechnology) and dorsal (9 min post-injection) and ventral (10.5 min post-injection) images were captured. To assess bioluminescence rhythms, anesthesia, D-luciferin injection and imaging was repeated at 4- to 8-hour intervals over approximately 30 hours. Similarly, female kidney reporter mice (*Ksp1.3-Cre; Dbp^KI/+^*) were imaged in a separate experiment to assess rhythmicity in bioluminescence, with 5 time-points distributed over 48 h. IVIS images were analyzed using Caliper Life Sciences’ Living Image software (version 4.4) within Regions of Interest (ROI) of fixed size.

Whole-body reporters (*Dbp^Luc /+^*) and liver reporters (*Alb-Cre+;Dbp ^KI/+^*) were also used to assess the distribution of bioluminescence by IVIS imaging. Mice were anesthetized with isoflurane, shaved, and injected with D-luciferin (100 microliters at 7-10 mM, i.p.) at times of peak expression (ZT 11-16). Images were captured of ventral and dorsal views at 9-12 minutes after injection. Bioluminescent counts within regions of interest (ROIs) were calculated using Living Image software. ROIs identified on the ventral surface were the whole rectangular region containing the mouse, and sub-ROI’s were a region in the throat (submandibular gland), upper abdomen, and lower abdomen. Dorsal ROI’s were the rectangle containing the entire mouse and a sub-ROI over the lower back, corresponding to the abdomen on the dorsal side. Subsequent calculations were performed in Microsoft Excel.

Liver and kidney reporter mice were anesthetized, dissected and imaged to confirm that bioluminescence originated exclusively from the liver. In additional animals, animals were euthanized before image collection.

### Bioluminescence Imaging of SCN Explants

Coronal sections containing SCN from adult *NMS-Cre;Dbp^KI/+^*, *AVP-IRES-CRE;Dbp^KI/+^*, and *Dbp^Luc/+^* mice were dissected, cultured, and imaged as previously described (Evans et al., 2011; Evans et al., 2013). Briefly, sections containing SCN (150 μm) were cultured on a Millicell membrane in air-buffered media containing 100 μM D-luciferin (Gold Biotechnology) and imaged for 5 days using a Stanford Photonics XR/MEGA-10Z cooled intensified charge-coupled device camera.

Rhythmic parameters of luciferase expression were calculated for each slice and for cell-like regions of interest (ROIs) within each slice using computational analyses in MATLAB (R2018a, MathWorks) as described previously (Evans et al., 2013; Leise & Harrington, 2011). Briefly, to locate and extract data from cell-like ROIs, we employed an iterative process identifying clusters of at least 20 bright pixels after background and local noise subtraction (through application of a 2D wavelet transform using Wavelab 850, (https://statweb.stanford.edu/~wavelab/) of a slice image summed across 24 h of bioluminescence. To extract time series for the ROI’s, each image in the sequence was smoothed via convolution with a Gaussian kernel applied to 12×12-pixel regions and reduced from 512×640 resolution to 256×320. A discrete wavelet transform (DWT) was applied to each time series to remove the trend and to extract the circadian and noise components using the *wmtsa* toolbox for MATLAB (https://atmos.uw.edu/~wmtsa/). The criteria for circadian rhythmicity in the ROI time series were a peak autocorrelation coefficient of at least 0.2, a circadian component peak-to-peak time between 18 and 30 h, an amplitude above baseline noise (standard deviation of noise component), and a cross-correlation coefficient of at least 0.4 with an aligned sine wave over a 48h window. Peaks of the DWT circadian component were used to estimate peak time of each ROI.

Rhythmicity index (RI) is the peak in the autocorrelation of the DWT-detrended time series, corresponding to a lag between 16 and 36 hrs, as previously described (Leise et al., 2013; Leise, 2017). The time of peak bioluminescence, rhythmicity index and the scatter of peak times within each slice for each ROI was assessed on the first day *ex vivo*. Period of rhythmicity in each ROI was determined as the average peak-to-peak interval in the second and third cycles. These measures were compared between genotypes by a general linear model, with slice ID included as a random variable to account for multiple cells being measured on each slice. Where applicable, post-hoc comparisons were performed using Tukey’s HSD pairwise comparisons.

### Data Collection and Analysis of Bioluminescence Rhythms in Ambulatory Liver Reporter Mice

Bioluminescence was measured in freely moving *Alb-Cre^+^; Dbp^KI/+^* reporter mice with the “Lumicycle *In Vivo*” system (Actimetrics, Wilmette, IL) using methods as recently described (Martin-Burgos et al., 2020). Animals were checked daily at varied times using an infrared viewer (Carson OPMOD DNV 1.0), or goggles (Pulsar Edge Night Vision Goggles PL75095).

Each Lumicycle *In Vivo* unit contained two PMTs (Hamamatsu H8259-01), and programmable LED lights. A programmable shutter blocked the PMTs during periods of light exposure and to measure ‘dark counts’. Each 1-minute dark-count value was subtracted from the counts recorded during the subsequent 14 minutes to obtain the background-corrected count values, to compensate for the effect of temperature fluctuations on PMT signal.

Ambulatory bioluminescence data were analyzed using RStudio. A discrete wavelet transform (DWT) was applied to each time series to detrend and to calculate the time of peaks using the wmtsa R package (https://cran.r-project.org/web/packages/wmtsa/index.html), as described (Leise & Harrington, 2011; Leise et al., 2013; Leise, 2017). The S12 filter was applied on 15-min median binned data; medians were used (instead of means) to reduce the effect of large outliers. Data before the first trough and after the last trough were discarded to avoid edge effects.

Locomotor activity was recorded using passive infrared motion sensors (Visonic, K940) and Clocklab software (RRID:SCR_014309). The mid-point of locomotor activity was determined by wavelet analysis on each day of recording. Midpoints were used because the onset of locomotor activity is poorly defined using motion sensors (relative to running wheel onsets).

### Assessing Routes of Administration of Luciferin

To determine whether rhythmic substrate intake influences the pattern of bioluminescence, we compared the time of peak bioluminescence between animals receiving continuous administration of substrate (from a subcutaneous osmotic minipump) with trials in which mice received D-luciferin in the drinking water (2 mM) and implantation of a PBS-filled osmotic pump.

Liver reporter mice previously housed in 12L:12D were entrained to a skeleton photoperiod (SPP) consisting of four 1-hour light pulses. A skeleton photoperiod provides additional periods of darkness in which to record bioluminescence. The use of a 4-pulse SPP (rather than the more typical 2-pulse SPP) was based on preliminary studies indicating a 4-pulse SPP could more consistently cause phase advances of locomotor activity following an advance shift of the lighting cycle. In this 4-pulse SPP, illumination occurred in four 1-hour blocks within the light phase in the preceding lighting cycle (e.g., lights were on from ZT 0-1, 2-3, 9-10, and 11-12, so the first and last hours of light in SPP coincided with the first and last hours of illumination in the full photocycle (with lights on ZT0-12 and lights off ZT12-24/0).

On the seventh day of SPP entrainment, mice were given analgesics (0.05 mg/kg Buprenorphine and 2.0 mg/kg Meloxicam), anesthetized with 3% isoflurane, shaved from hips to shoulders, and a primed osmotic minipump (Alzet Model #1002, 0.25µl per hour, 14 day) containing D-luciferin (100 mM dissolved in PBS) or PBS vehicle was implanted subcutaneously. Mice were returned to their cages with a warming disc and were provided soft food during the first 24 hours of recovery. Animals were placed into the LumiCycle *In Vivo* unit 2.5 days after surgery. Bioluminescence was recorded in SPP lighting for 2.5 days, then lights were disabled at the time of lights-out. The time of peak bioluminescence was determined by wavelet analysis on the first day in constant darkness. No difference in peak time of bioluminescence was found (see Results); in subsequent studies of ambulatory Liver reporter mice, D - luciferin (2 mM) was administered in the drinking water.

### Re-entrainment following a Phase Shift of the Skeleton Photoperiod

Liver reporter mice (*Albumin-Cre; Dbp^KI/+^*) previously entrained to LD were transferred to the skeleton photoperiod for several days. Mice were anesthetized with isoflurane and shaved 2.5 days prior to placement in the LumiCycle *In Vivo* units. D-Luciferin (2 mM) was provided in the drinking water. Skeleton photoperiod lighting conditions were either maintained at the initial pattern or advanced by 6 hr after the second day of recording. Locomotor activity was detected by passive infrared motion sensors.

The circadian time of peak bioluminescence and the mid-point of locomotor activity were determined by wavelet analysis on each day of recording. We used the midpoint of locomotor activity because activity onset was not easily defined using motion sensors. The timing of bioluminescence and locomotor activity rhythms was normalized to the timing of these rhythms on Day 2 (e.g., the last day before shifting the lighting cycle in the shifted group) for each animal. Data are expressed as mean ± SEM for each lighting condition and endpoint on each day. Data from each lighting group were analyzed separately using a general linear model with Animal ID as a random variable (allowing comparison of the two rhythms within individuals) and the main effects of the endpoint measure (locomotor activity or bioluminescence) and Day number, and the 2-way interaction Measure*Day. In animals not undergoing a phase shift, potential changes in the timing of the locomotor or bioluminescence rhythm were assessed separately for either measure by testing the influence of Day number.

### Food Restriction Followed by Bioluminescence Recording

Liver reporter mice (*Albumin-Cre; Dbp^KI/+^*) were fed pellets (300 mg, Dustless Precision Pellets, Rodent, Grain-Based, F0170, BioServ, Flemington, NJ, USA) through the Actimetrics timed feeding apparatus designed by Phenome Technologies, Skokie, IL, USA. Pellets were spaced by a minimum of 10 minutes to prevent hoarding behaviour (Acosta-Rodriguez et al., 2017). Liver reporter animals were randomly assigned to treatment groups and recording boxes. Three groups were studied: those with *ad libitum* access to food, those with feeding restricted to the light phase of the LD cycle (daytime feeding), and mice with access to food restricted to the dark phase of the LD cycle (nighttime feeding). Mice were weighed regularly to ensure body weight did not decrease below 95% of initial weight. All mice were kept on a 12L:12D lighting schedule during the period of food manipulation, and then were released into constant darkness with D-luciferin (2mM) in the drinking water for bioluminescence recording. During the LD period, data were collected on feeding, light levels, and locomotor behavior (using motion sensors). Three days before entering the LumiCycle *In Vivo* units, cage bottoms were changed at dark onset. *Ad libitum* and night-fed mice were placed into the LumiCycle *In Vivo* units at dark onset with food immediately available. Day-fed mice were placed into the LumiCycle *In Vivo* units at dark onset but were provided food after 12 hours (at the time of light onset in the previous LD cycle) to continue the daytime feeding regime during the first day of the recording period. Bioluminescence was recorded for 7 days.

Experimental groups and controls ran in parallel over five cohorts lasting 3 months. 24 hours prior to placement in the recording boxes, mice were shaved from hips to shoulders on their front and back under 3% isoflurane and returned to their cages. Mice were provided with D-luciferin (2mM) in the drinking water 6 hours prior to placement into the LumiCycle *In Vivo* units, to enable instantaneous bioluminescence upon recording onset.

The center of gravity (COG) of food intake was calculated for each animal for the last 5 days in LD (e.g., the last 5 days of the feeding regimen). Food intake patterns were also independently assessed qualitatively by four observers. These assessments led to identification of three cohorts of mice, based on food intake patterns. Three mice were identified as clear outliers compared to these three cohorts based on visual inspection of the food intake timing. In line with this qualitative assessment, the feeding COG of each of these 3 animals was >2 h removed from the other animals in their cohort. These three animals were excluded from cohort-based assessments. Peak of bioluminescence on each day was calculated by DWT analysis as above. Missing data resulted from inability to define a time of peak bioluminescence on some days. Hair regrowth contributed to loss of signal and loss of rhythm amplitude, and thus to missing data in some cases.

### Data and Materials Availability

Requests for research materials should be directed to Dr. David Weaver. Underlying data are available from Dr. Weaver on request.

## Results

### Generation of a bifunctional reporter mouse

CRISPR/Cas9 genome editing was used to introduce a bifunctional reporter into the mouse *Dbp* locus (**Fig. 1**). The reporter consists of a T2A sequence (to allow expression of separate proteins from a single transcript), a destabilized, enhanced GFP (d2EGFP, hereafter GFP) sequence flanked by loxP sites, and a codon optimised synthetic firefly *luciferase* (*Luc2* from *Photinus pyralis,* hereafter luc). In the absence of *Cre* expression, DBP and GFP are expressed as separate proteins. After CRE-mediated recombination, the floxed GFP is removed, and separate DBP and luciferase proteins are expressed from the *Dbp* locus. Sequencing of genomic DNA confirmed successful generation of the *Dbp^KI^* conditional reporter allele.

A non-conditional reporter allele was generated by breeding to combine the conditional *Dbp^KI^* allele with *Cre*-recombinase expressed in the germline (of a female *Prrx1-Cre* mouse), leading to germline excision of GFP. We refer to this non-conditional allele, which expresses luciferase wherever *Dbp* is expressed, as *Dbp^Luc^*.

### Molecular and Behavioral Rhythms in Mice with *Dbp* Reporter Alleles

To confirm that the introduction of the reporter construct into the *Dbp* locus did not alter circadian clock function, molecular and behavioral rhythms were assessed. Mice used for these analyses had either one or two copies of the GFP-containing conditional allele (*Dbp^KI/+^* and *Dbp^KI/KI^*, respectively), one or two copies of the luciferase-expressing allele (*Dbp^Luc/+^* and *Dbp^Luc/Luc^*, respectively), or were wild-type (WT) littermate controls.

RNA was isolated from male livers collected at 4-h intervals over 24-h in a 12L:12D (LD) lighting cycle. Northern blots were prepared and probed for *Dbp* and *Actin* (loading control). As expected, the transcripts from *Dbp^KI^* and *Dbp^Luc^* alleles migrated more slowly than the wild-type transcript (**Fig. 2A**), due to inclusion of GFP and luciferase coding sequence in these transcripts, respectively, as verified by probing for reporter sequences in a separate blot. Peak levels of *Dbp* expression in liver occurred at ZT10 in all genotypes (**Fig. 2B, 2C**), as expected based on previous studies^3,42,57^. For each transcript type, the *Dbp*/*Actin* ratios were ranked within each series of 6 timepoints. These ranks differed significantly among the timepoints for each transcript (Friedman’s One-Way analysis of variance, *p* < 0.002), and post-hoc testing indicated significantly higher rankings at ZT10 than at ZT2, ZT18 and ZT22 (Dunn’s test, *p* < 0.05; Fig 2D-2F). These data indicate that the temporal profile of transcript expression from the *Dbp* locus was unaffected by the inclusion of reporter sequences.

**Figure 2.**
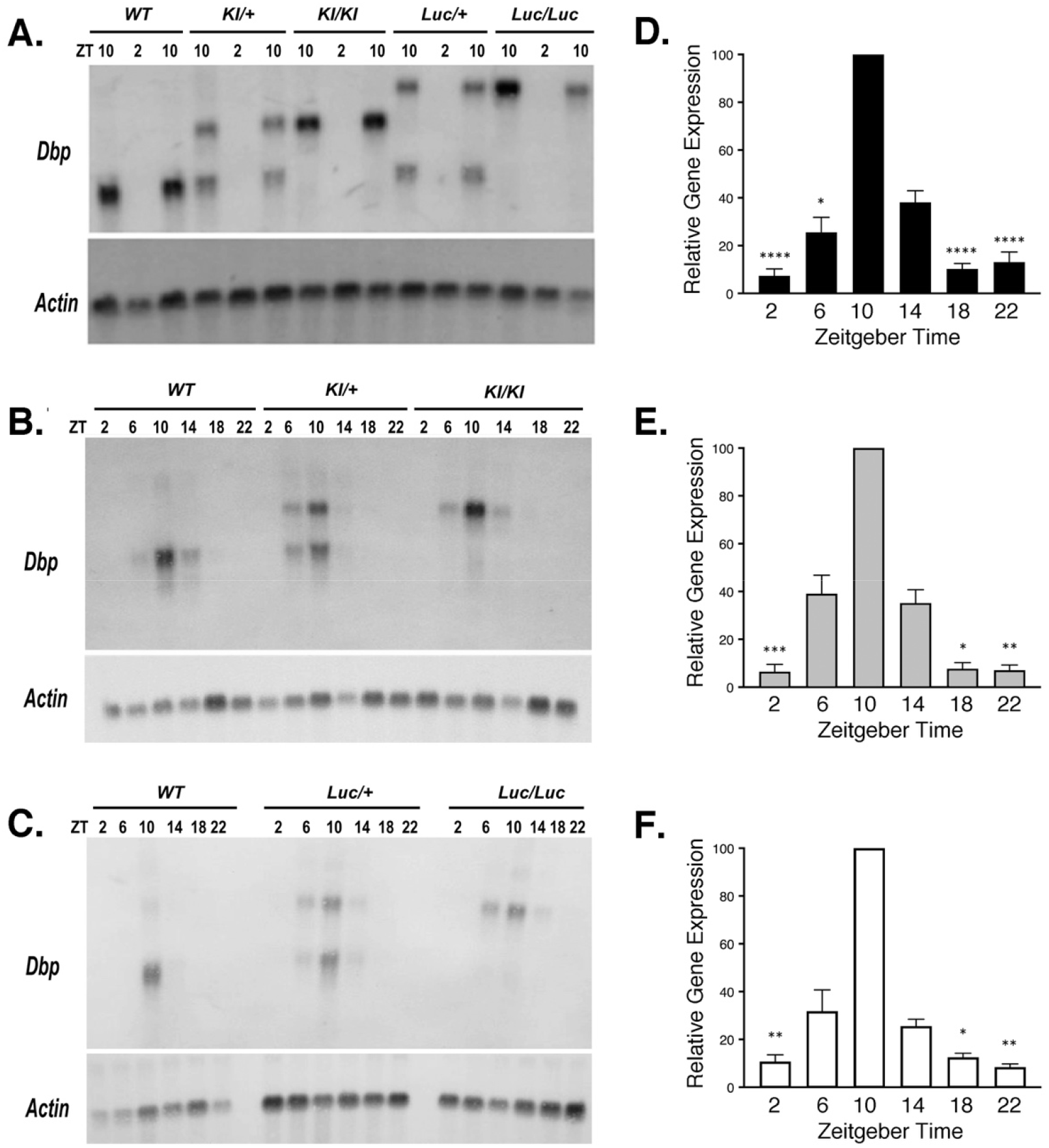
*Dbp* mRNA rhythms are not altered in reporter mice. **A-C**. Representative Northern Blots probed to detect *Dbp* and *Actin* mRNA. **A.** From each of five genotypes, RNA samples were extracted from livers collected at ZT 2 and 10. For each genotype, there are two samples at ZT10 and one sample at ZT2 on this blot. **B**. and **C**. Representative Northern Blots of RNA samples collected from WT and reporter mouse livers at each of six Zeitgeber times (ZT). **D-F**. Quantification of *Dbp* mRNA rhythms for each allele in time-series experiments (6 time-points each). Results are expressed as mean (± SEM) percent of the peak *Dbp*/*Actin* ratio, which occurred at ZT 10 on every blot. **D.** Wild-type *Dbp* transcript (n=12 sample sets). **E.** *Dbp^KI^* transcript (n= 6). **F.** *Dbp^Luc^* transcript (n= 6). For each transcript, there was a significant rhythm (Friedman’s One-way ANOVA, Q > 19, *p* < 0.002). Asterisks indicate time-points that differed significantly from ZT10 (Dunn’s test, * *p* < 0.05, ** *p* < 0.01, *** *p* < 0.001, **** *p* ≤ 0.0001). Significant differences among some other time-points are not shown for clarity.

Heterozygous mice expressed both *Dbp* and *Dbp-plus*-*reporter* transcripts. The two transcript types did not differ in abundance: optical density over film background of the *Dbp^KI^* transcript was 100.5 ± 5.3 % of the *Dbp^+^* transcript in *Dbp^KI/+^* mice (t=0.084, df=7, p= 0.94, one-sample t-test vs 100%), while the *Dbp^Luc^* transcript was 102.3 ± 5.0 % of *Dbp^+^* transcript in *Dbp^Luc/+^* mice (t=0.446, df=7, p=0.669). The equivalent expression level of the two transcript types in heterozygous animals strongly suggests that transcript regulation and stability were not altered by inclusion of reporter-encoding sequences.

Locomotor activity rhythms were assessed in constant darkness in mice of both sexes in the same five genotypes (**Table 1; Fig. S1**). We found a significant sex-by-genotype interaction (F_4,102_ = 2.904, *p* = 0.0254). Post-hoc tests indicated an unexpected sex difference in the *Dbp^Luc/Luc^* mice. Indeed, when this genotype was excluded from the analysis, no significant sex-by-genotype interaction was observed (F_3,88_ = 1.349; *p* = 0.2636) and one-way ANOVA did not find a significant main effect of genotype (F_3,91_ = 1.174; *p* = 0.3242). One-way ANOVA within each sex with all five genotypes included revealed no genotype effect in males (F_4,50_ = 1.299, *p* = 0.283). While there was a significant genotype effect in females (F_4,52_ = 2.716, *p* = 0.040), Tukey HSD post-hoc tests did not find a significant result among any of the pairwise genotype comparisons (all p values > 0.05). Similarly, an alternative post-hoc analysis revealed that none of the other female genotypes differed from WT females in their free-running period in constant darkness (Dunnett’s test, p > 0.5 in each case). To further examine the effect of sex on free-running period, males and females of each genotype were compared directly. In both *Dbp^Luc/Luc^* and *Dbp^KI/KI^* mice, males had significantly longer periods than females (*p* < 0.01), while there was no sex difference in wild-types or heterozygous reporters (*p* > 0.46).

**Table 1:** Period length of locomotor activity rhythms in constant darkness, by sex and genotype

| Genotype | Sex | N | $\tau_{DD}$ (Mean ± SEM), h |
| --- | --- | --- | --- |
| *Dbp*<sup>+/+</sup> | Male | 15 | 23.88 ± 0.027 |
| *Dbp*<sup>KI/+</sup> | Male | 10 | 23.91 ± 0.057 |
| *Dbp*<sup>KI/KI</sup> | Male | 11 | 23.92 ± 0.036 |
| *Dbp*<sup>Luc/+</sup> | Male | 11 | 23.86 ± 0.025 |
| *Dbp*<sup>Luc/Luc</sup> | Male | 8 | 23.97 ± 0.029 |
| *Dbp*<sup>+/+</sup> | Female | 21 | 23.87 ± 0.021 |
| *Dbp*<sup>KI/+</sup> | Female | 9 | 23.89 ± 0.036 |
| *Dbp*<sup>KI/KI</sup> | Female | 11 | 23.79 ± 0.030 |
| *Dbp*<sup>Luc/+</sup> | Female | 8 | 23.82 ± 0.053 |
| *Dbp*<sup>Luc/Luc</sup> | Female | 8 | 23.75 ± 0.042 |

Together, these assessments of molecular and behavioral rhythms indicate that the reporter alleles do not change *Dbp* expression or appreciably alter circadian function.

### GFP expression from the *Dbp^KI^* allele

To examine expression of GFP from the conditional allele, *Dbp^KI^*^/+^ mice (n=5-6 mice per time-point) were anesthetized and perfused with fixative at 4-h intervals over 24 h (**Fig. S2**). Liver sections from *Dbp^KI/+^* and control (WT) mice were examined by confocal microscopy. Fluorescence signal intensity did not differ between time-points (ANOVA F_5,26_ =1.279, *p* = 0.7560). GFP signal from *Dbp^KI/+^* liver sections was 5-10x higher than from WT sections, but absolute levels were quite low. The low level of GFP expression may be due to the use of destabilized GFP with a 2-hour half-life, intended to more accurately track changes on a circadian time-scale. The relatively low level and lack of detectable rhythmicity in GFP expression was unexpected, especially considering that liver is the tissue with the highest levels of *Dbp* expression (Fonjallaz et al., 1996) and thus may represent a ‘best-case’ scenario. As the primary objective of this project was to generate a mouse model with *Cre*-dependent expression of bioluminescence from the *Dbp* locus, however, the absence of robust GFP-driven fluorescence rhythms in *Cre*-negative cells did not preclude achieving this objective. GFP is effectively serving as a ‘floxed stop’ to make luciferase expression from the *Dbp* locus exclusively *Cre*-dependent.

### Non-conditional luciferase expression from the *Dbp^Luc^* allele

The *Dbp^Luc^* allele produces widespread, rhythmic luciferase expression, both *in vivo* and *ex vivo*. More specifically, explants of lung and anterior pituitary gland from *Dbp^Luc/+^* mice incubated with D-luciferin had robust circadian rhythms in bioluminescence (**Fig. 3**). Furthermore, *in vivo* imaging of *Dbp^Luc/+^* mice at 7 time-points over a ∼30-h period revealed rhythmic bioluminescence in the abdomen and throat in ventral views, and in the lower back in dorsal views (**Fig. 4B**), similar to the distribution of bioluminescence signal from *Per2^Luciferase/+^* (Tahara et al., 2012) and *Per2^LucSV/+^* mice (van der Vinne et al., 2018; van der Vinne et al., 2020) (**Fig. 4A**). The level of light output was ∼2.5-fold greater in ventral views than in dorsal views (p<0.0001, Wilcoxon matched pairs test, W=151, n=17). In the abdomen, a rostral (“liver”) region of interest (ROI) accounted for 46.6 ± 3.0% (Mean ± SEM; n=17) of bioluminescence from the ventral view, while the lower abdomen contributed another 38.4 ± 3.5%. Bioluminescence rhythms from the throat region of *Per2^Luciferase^* mice have previously been shown to originate in the submandibular gland (Tahara et al., 2012). Bioluminescence was absent in mice with wild-type *Dbp* alleles or with the conditional *Dbp^KI^* allele (in the absence of *Cre*).

**Figure 3.**
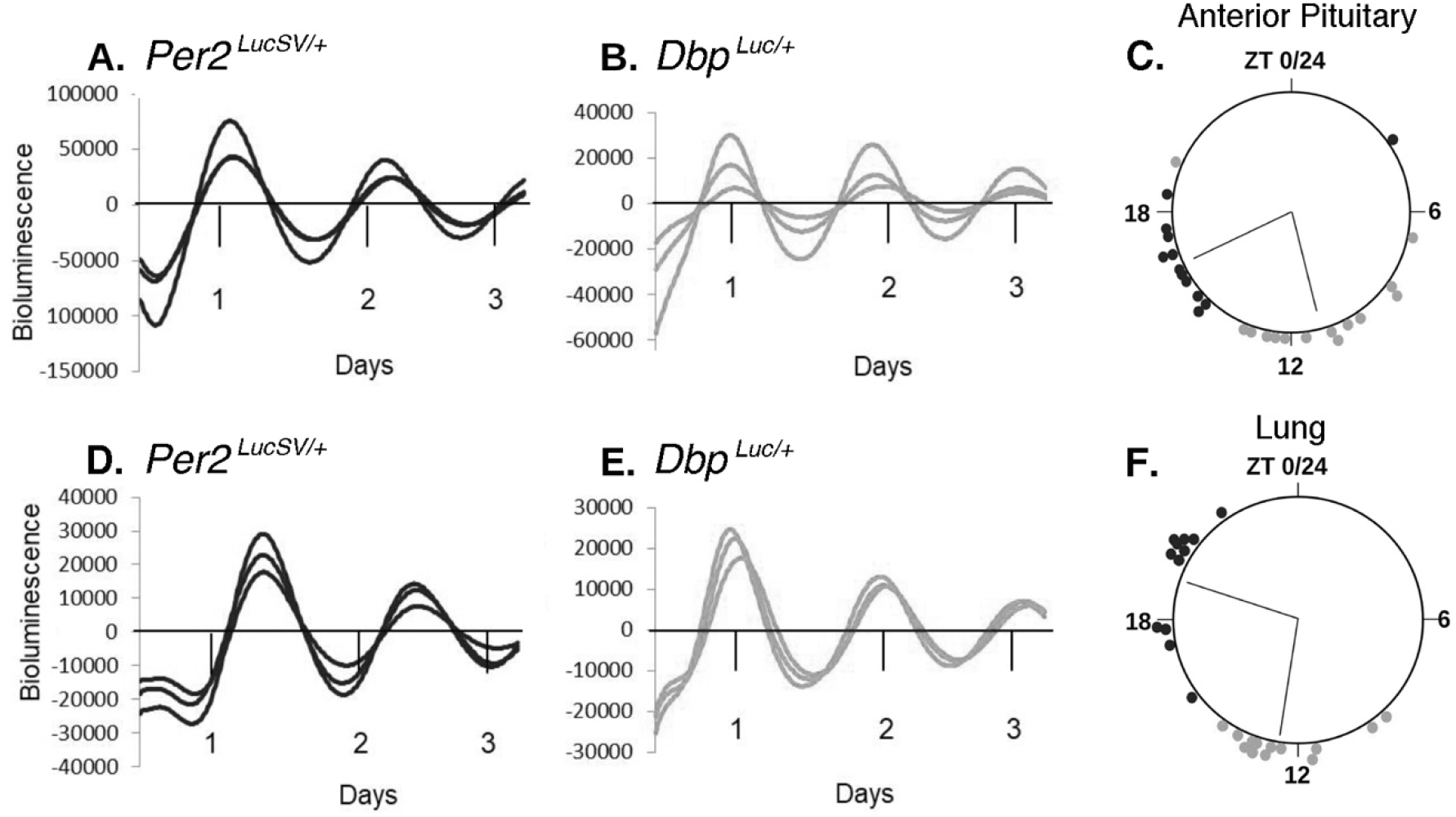
*Ex vivo* bioluminescence rhythms from *Per2^LucSV/+^* and *Dbp^Luc/+^* tissue explants. **A-C.**, Anterior Pituitary gland. **D-F.**, Lung. **A., B., D., and E.** are representative bioluminescence rhythms from triplicate tissue explants from *Per2^LucSV/+^* (**A., D.**) and *Dbp^Luc/+^* mice (**B., E.**). ‘Days’ refers to time in culture, not projected ZT. Values are 24-h background-subtracted and 3-h smoothed. **C., F.,** Time of peak bioluminescence *ex vivo.* The large circles represent a 24-h day for each organ. ZT’s refer to the lighting cycle to which the mice were exposed prior to sample collection, with ZT0-12 being the light phase. Circles at the perimeter of the large circle indicate the timing of peak bioluminescence of individual *Per2^LucSV/+^* (black) or *Dbp^Luc/+^* (gray) tissue explants (n=12-14 mice). Within each tissue/genotype combination, there was significant clustering of times of peak bioluminescence. Radial lines represent the mean peak time, which differed significantly between genotypes for each tissue (Watson-Williams test, p<0.001).

**Figure 4.**
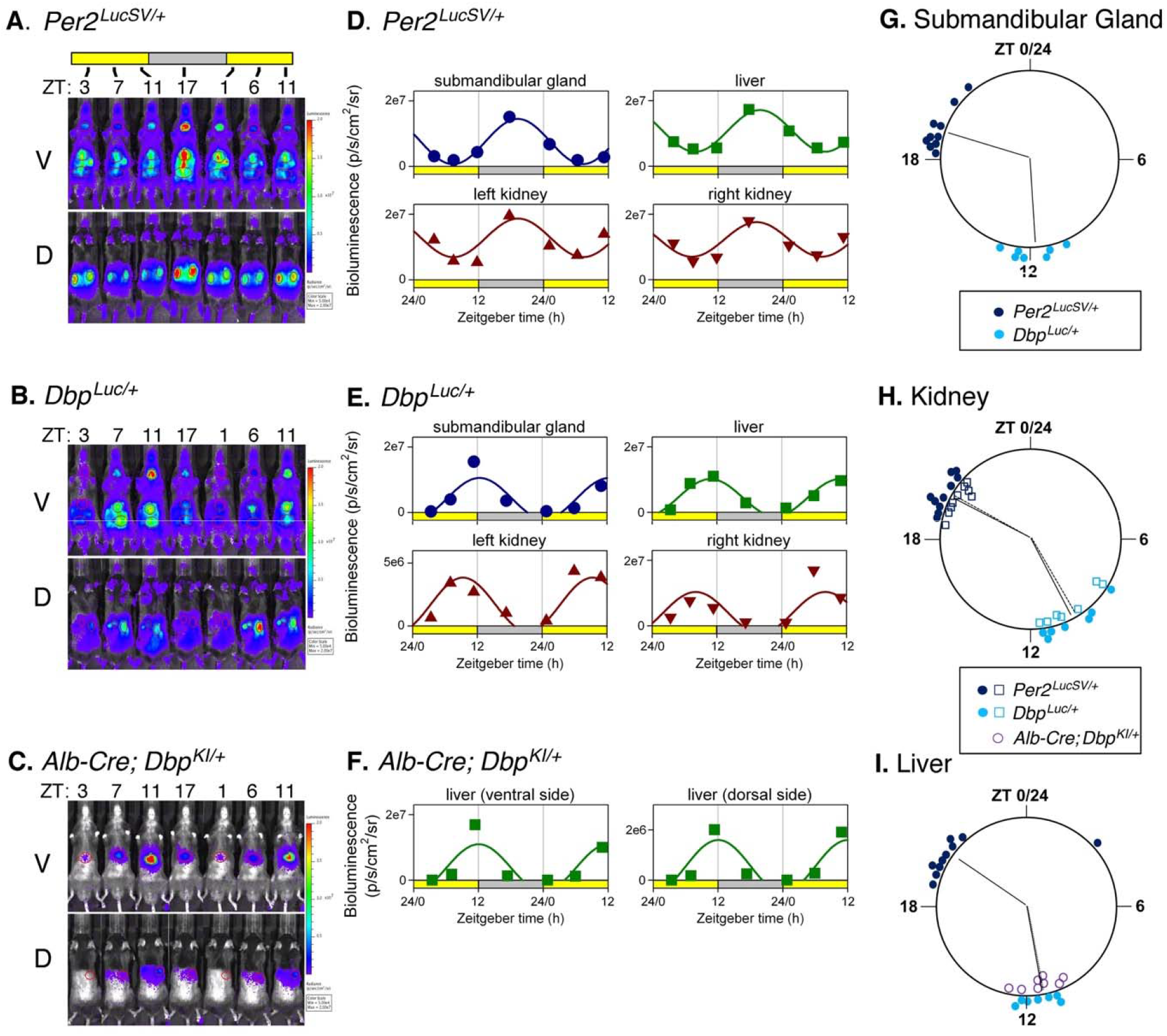
Bioluminescence rhythms measured *in vivo*. **A-C**. Bioluminescence images captured at 4-6 hr intervals from a representative mouse of each genotype. **A.** *Per2^LucSV/+^*, **B**. *Dbp^Luc/+^* **C**. *Alb-Cre+ ; Dbp^KI/+^.* Ventral (V) and dorsal (D) views are shown for each mouse. All images for each mouse are set to the same luminescence scale. **D-F**. Cosinor-fitting of bioluminescence signal over time for the animals shown in Panels A-C to determine peak time. Bioluminescence rhythms were assessed in submandibular gland, liver, and kidneys of (**D.**) *Per2^LucSV/+^* and (**E.**) *Dbp^Luc/+^* reporter mice, and from liver of *Alb-Cre+ ; Dbp^KI/+^* mice (**F.**). **G-I.** Time of peak bioluminescence *in vivo*. **G.** Submandibular gland, **H.** Kidneys, and **I.** Liver. Data plotted as in Fig. 3. *Per2^LucSV/+^* tissues (n=10, dark blue), *Dbp^Luc/+^* tissues (n=7, teal). In Panel H, open squares and filled circles represent the right and left kidneys, respectively. In Panel I, purple circles represent livers from *Alb-Cre+ ; Dbp^KI/+^* mice (n=8). Radial lines represent the mean peak time for each genotype and tissue. Radial lines from the two kidneys of a genotype are nearly overlapping. For liver, radial lines for the two *Dbp* reporter lines are overlapping and appear as a single line. Within each organ examined, time of peak differed significantly in *Per2^LucSV/+^* explants compared to *Dbp^Luc/+^* and *Alb-Cre+ ; Dbp^KI/+^* explants (*p*=0.002, Watson-Williams test). There was no significant difference in peak time between *Dbp^Luc/+^* and *Alb-Cre+ ; Dbp^KI/+^* liver tissues (*p*>0.05).

Previous reports have shown that in a number of tissues, *Dbp* RNA levels peak earlier than *Per2* RNA levels (Punia et al., 2012; Zhang et al., 2014). Consistent with this literature, the time of peak of bioluminescence rhythms from *Dbp^Luc/+^* tissues preceded the time of peak of bioluminescence rhythms from *Per2^LucSV/+^* tissues by ∼ 6 hours in explants (**Fig. 3C, 3F**) and by ∼ 9 hr *in vivo* (**Fig. 4G-4I).** Bioluminescence rhythms from *Per2^LucSV/+^* tissue explants had significantly longer period than explants from *Dbp^Luc/+^* mice (Lung: 25.29 ± 0.13 vs 23.93 ± 0.11 h; *F*_1,27.7_ = 95.55, *p* < 0.0001; Anterior Pituitary: 25.27 ± 0.08 vs 23.73 ± 0.112 h; *F*_1,24.53_ = 66.12, *p* < 0.0001).

### *Cre*-dependent Luciferase Expression in Liver

The main use we envision for the *Dbp* reporter alleles involve *Cre* recombinase-mediated excision of GFP, leading to expression of *luciferase* in cells expressing *Cre*. The effectiveness of this approach was first assessed in hepatocytes using an *Albumin-Cre*-driver line. *In vivo* bioluminescence imaging of intact *Albumin*-*Cre*^+^; *Dbp^KI^*^/+^ “liver reporter” mice at the time of expected maximal bioluminescence revealed that 96.6 ± 0.48% of light originated in the “liver” ROI (relative to total ventral-view bioluminescence; p<0.0001 versus 46.6 ± 3.0% in *Dbp^Luc^* mice, U-test, U=0, n=19 and 17, respectively). Notably, post-mortem imaging after dissection confirmed that bioluminescence originated exclusively from the liver in these mice (97.4% of light from liver; n=12).

In a separate cohort of liver reporter mice, bioluminescence was assessed around the clock by IVIS imaging. The cosinor-fitted time of peak of *Dbp*-driven bioluminescence rhythms from the liver ‘region of interest’ of these mice (ZT11) was indistinguishable from the peak time of the liver ROI analyzed in whole-body *Dbp^Luc^* mice (**Fig. 4I**).

### *Cre*-dependent Luciferase Expression in Kidney

Viral introduction of rhythmic luciferase reporters to the liver has been used previously (Saini et al., 2013, Sinturel et al., 2021), so our success with detecting bioluminescence rhythms specifically from the liver in *Albumin*-*Cre*^+^; *Dbp^KI^*^/+^ “liver reporter” mice was reassuring, but not surprising. With reporter genes expressing from multiple tissues (e.g., *Dbp^Luc^* and *Per2^Luc^*), the contribution made by surrounding organs may be unclear. To extend our demonstration of tissue-specific luciferase expression from the conditional *Dbp^KI^* allele, we examined bioluminescence from anesthetized *Ksp1.3-Cre; Dbp^KI^*^/+^ “kidney reporter” mice. The *Ksp1.3-Cre* driver leads to recombination in the developing kidney and urogenital tissues, and in renal tubules of adult mice. In male kidney reporter mice, IVIS imaging of anesthetized, dissected living mice revealed bioluminescence from the kidney and seminal vesicles *in situ* (**Fig. S3, S4**). In females, bioluminescence originated from the kidney and proximal ureter (**Fig. S5**). We thus used female mice to assess rhythmicity. Clear diurnal rhythmicity in bioluminescence was apparent from the kidney (Friedman’s One-Way Analysis of Variance, (F_r_ =32.71, k=5, n=9, p<0.0001, see **Fig. S6**), with a peak at ZT8. Dunn’s test revealed that ZT8 timepoint differed significantly from ZT0 and ZT18 but not from ZT4 and ZT14 (multiplicity-corrected, two-tailed Dunn’s test; see **Fig. S6**).

### Cell-type Specific Bioluminescence Rhythms in SCN Slices

The heterogeneity of SCN neurons has important functional implications for our understanding of the central circadian clock (Herzog et al., 2017). Neuromedin S (NMS) is expressed in ∼40% of SCN cells, while Arginine Vasopressin (AVP) is expressed in ∼10% of SCN neurons and is contained within the NMS-expressing population (Lee et al., 2015). The utility of our conditional reporter line was demonstrated by monitoring bioluminescence rhythms within specific subpopulations of SCN neurons (**Fig. 5**). *NMS-iCre*; *Dbp^KI/+^* mice and *AVP-IRES2-Cre; Dbp^KI/+^* mice were generated, and single-cell bioluminescence rhythms were compared to those from non-conditional *Dbp^Luc/+^* mice in SCN slices *ex vivo*. For the conditional mice, bioluminescence was apparent in subsets of cells within the SCN (**Fig. 5A**). The anatomical pattern of bioluminescence in the SCN differed based on the *Cre* line used, consistent with the expected distribution for each neuronal subtype. In each slice, rhythmic ROI’s were readily apparent (**Fig. 5B**).

**Figure 5.**
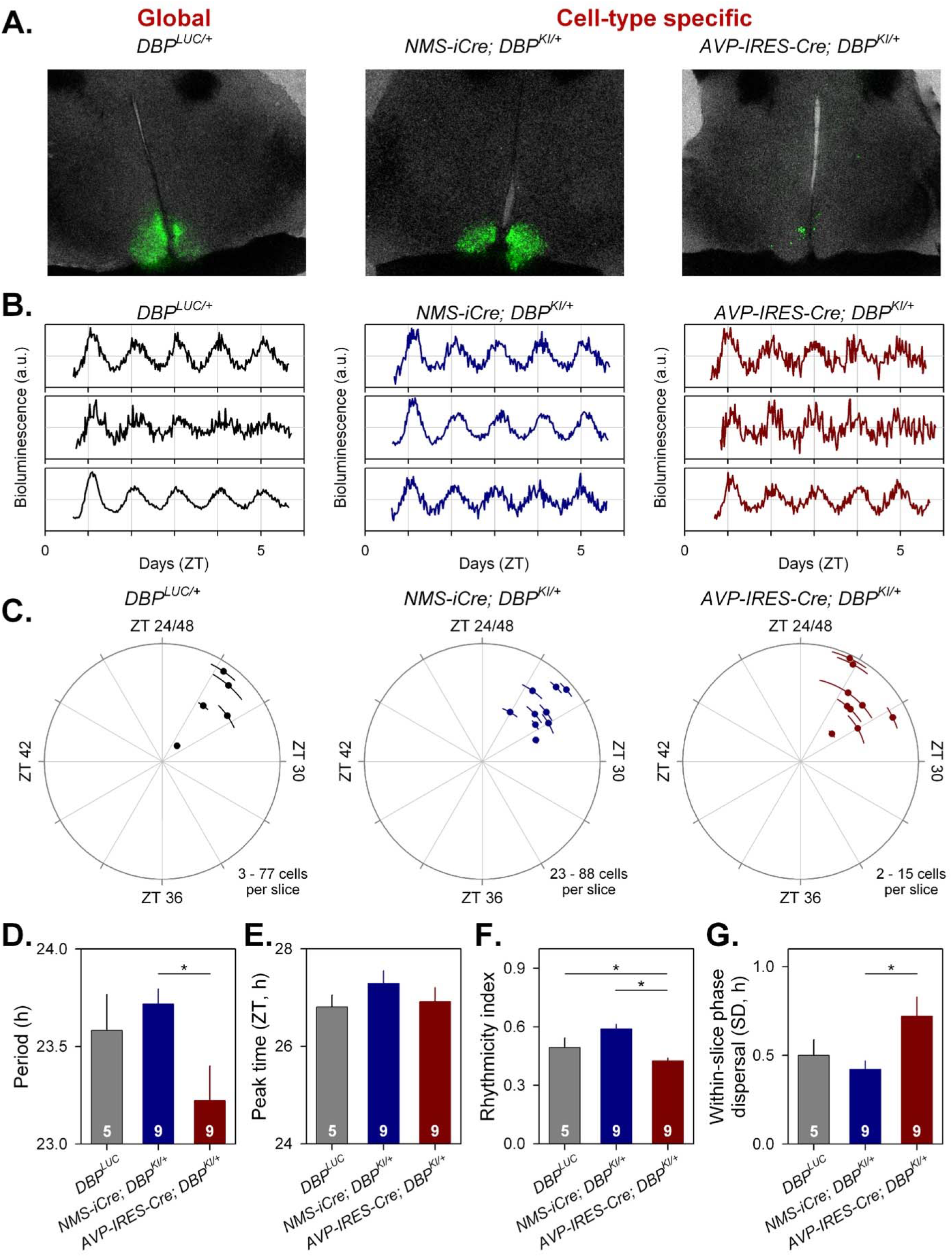
Cell-type-specific imaging of luciferase expression in SCN slices. **A**) 24h summed bioluminescence overlaid onto bright field images of a section through the SCN from *Dbp^Luc/+^* (global reporter expression, left), and in mice expressing luciferase from specific subsets of SCN neurons (NMS^+^ cells, center; AVP^+^ cells, right). **B.** Representative bioluminescence traces from single neuron-like ROIs in slices from each genotype. **C.** Circular plots indicate the peak time of bioluminescence rhythms from each genotype. Time is expressed relative to the light-dark cycle the mice were housed in prior to sacrifice; numbers >24 are used to indicate that these measures are recorded on the first day in culture and are plotted relative to the previous lighting conditions. Each slice is represented by a small dot. Placement of the dot relative to outer circle indicates average peak time (±SD), while the distance from the center corresponds to the number of cells incorporated in the average (√cell#). **D-G**. Rhythm parameters by genotype. The number of slices per genotype is indicated at the base of each bar. **D.** Mean period (± SEM). **E.** Circular mean peak time (± SEM). **F.** Mean rhythmicity index score (± SEM). **G.** Mean peak time dispersal (quantified by circular SD of peak times within each slice).

The cell-type specificity of bioluminescence signals from the different genotypes enabled the assessment of rhythm quality in the different neural populations. This assessment revealed a significantly shorter period in AVP^+^ cells compared to NMS^+^ cells (**Fig. 5C, Fig. 5D**; F_2,14.64_ = 4.259, *p* = 0.0345). The time of peak of *Dbp*-driven bioluminescence did not differ significantly between the different cellular populations examined (**Fig. 5E**; F_2,18.31_ = 0.6570, *p* = 0.5302), while a reduction in rhythm robustness was observed in AVP^+^ neurons compared to rhythms of NMS^+^ neurons as well as compared to all cells (**Fig. 5F**; F_2,18.11_= 14.34, *p* = 0.0002). The distribution of peak times was also more dispersed in AVP^+^ cells compared to NMS^+^ cells (**Fig. 5G**).

These results complement the recent report from Shan *et al*. (2020) using a *Cre*-dependent Color-Switch PER2::LUC reporter mouse demonstrating period and phase differences among sub-populations of SCN neurons. Our *Dbp^KI^* mice and the recently reported Color-Switch PER2::LUC mouse line (Shan *et al*., 2020) will be important additions to our molecular-genetic armamentarium for unravelling the complicated relationships among the cellular components of the SCN circadian pacemaker^58-64^.

### Route of Substrate Administration

Monitoring peripheral organ circadian phase following disruptive environmental, surgical or genetic conditions will require long-term monitoring of peripheral rhythms in ambulatory mice. Studies using substrate delivery by mini-osmotic pump or infusion pump allow constant substrate administration but require surgery and, in the case of mini-osmotic pumps, are limited by the pump volume. Therefore, administration of luciferase substrate in the drinking water would be preferable. Thus, we examined the potential impact of route of substrate administration on rhythm phase using the Lumicycle *In Vivo* system (Actimetrics, Wilmette IL) in *Albumin-Cre*; *Dbp^KI^*^/+^ (“liver reporter”) mice. Mice were entrained to LD followed by a skeleton photoperiod consisting of four 1-h pulses of light every 24 hr (1L:1D:1L:6D:1L:1D:1L:12D) with the 12-h dark phase coinciding with 12-h dark phase of the preceding LD cycle. A skeleton photoperiod was used because detection of bioluminescence requires the absence of ambient light, while studies of light-induced phase shifting obviously require light; a skeleton photoperiod is a compromise between these conflicting constraints. After 7 days in the skeleton photoperiod, mice were anesthetized for subcutaneous implantation of a primed osmotic minipump (Alzet, Model #1002 (0.25µl per hour)) containing either D-luciferin (100 mM) or phosphate buffered saline (PBS). Mice with PBS-containing pumps received D-luciferin in the drinking water (2 mM). Rhythms of bioluminescence were readily detected under these conditions (**Fig. S7**). The time of peak bioluminescence was determined by discrete wavelet transform (DWT) analysis on the first day of exposure to constant darkness (5 days after pump implantation). There was no difference in time of peak between these routes of administration (drinking water: mean peak time (± SEM) CT 8.75 ± 0.20 (n = 7); osmotic minipumps: mean peak time CT 8.76 ± 0.19 (n=7); unpaired t-test, t = 0.0342, df =12, *p* = 0.9733). Thus, the presumed rhythm of substrate intake, secondary to the rhythm of water intake, does not change the time of peak of the bioluminescence rhythm. This is consistent with recent results from Sinturel et al., (2021) and Martin-Burgos et al., (2022) using PER2^Luciferase^ mice. Subsequent studies used D-luciferin (2 mM) administered in the drinking water.

### Circadian Misalignment Following a Phase Shift of the Lighting Cycle

The approach described above provides an unparalleled system for assessing the timing of rhythmicity in a specific tissue over long periods of time. Next, hepatic bioluminescence rhythms were monitored in *Albumin-Cre*; *Dbp^KI^*^/+^ (liver reporter) mice before and after a 6-hr phase advance of the skeleton lighting cycle. Mice that remained in the original (non-shifted) skeleton lighting regimen had a stable phase of hepatic bioluminescence (**Fig. 6C**). In contrast, mice exposed to a phase-advance of the skeleton photoperiod displayed a gradual phase-advance in both locomotor activity and hepatic bioluminescence rhythms (**Fig. 6A, B**). Notably, locomotor rhythms shifted more rapidly than hepatic bioluminescence (**Fig. 6B**). To compare the re-entrainment of bioluminescence and locomotor activity rhythms, peak time for each rhythm each day was normalized to the time of peak on the last day before shifting the lighting cycle in the shifted group (e.g., Day 2 in **Fig. 6**) for each animal. Data from each lighting group were analyzed separately using a general linear model with Animal ID as a random variable (allowing comparison of the two rhythms within individuals) and the main effects were endpoint (locomotor activity or bioluminescence) and Day number. In animals not undergoing a phase shift, the phase relationship of these endpoints was unchanged over time (F < 1.1, *p* > 0.39). In contrast, in animals exposed to a 6-hr phase advance, the phase relationship of the locomotor activity and bioluminescence rhythms differed significantly (Measure*Day interaction, F_9,54.98_ = 3.358, *p* = 0.0024). Post-hoc testing revealed a significant difference in phase between the two measures on day 9 (Tukey HSD, p<0.05). A separate analysis to compare phase (relative to Day 2 baseline) between bioluminescence and locomotor activity rhythms revealed significant differences between the two measures on days 5, 6, 7, 8, 9 and 10 (t-tests on each day, *p* < 0.05). Thus, both locomotor activity and hepatic bioluminescence rhythms shifted following a phase shift of the lighting cycle, but the rhythms differ in their kinetics of re-adjustment: liver lagged behind. These data provide clear evidence for misalignment of SCN-driven behavioral rhythms and hepatic rhythmicity.

**Figure 6.**
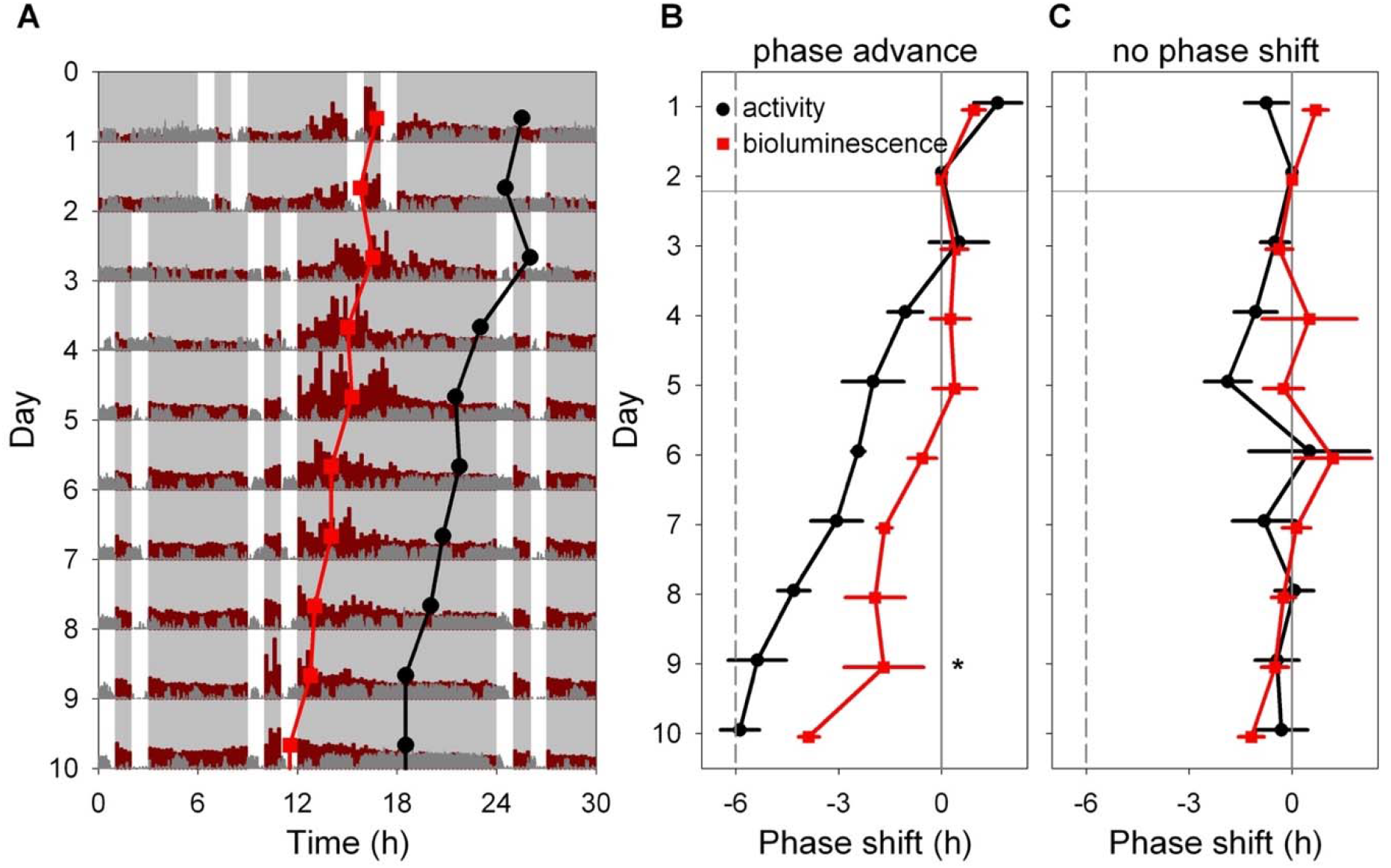
Light-induced resetting produces misalignment between rhythms in liver bioluminescence and locomotor activity. **A.** Representative double-plotted actogram showing locomotor activity (dark gray) and bioluminescence (dark red) of an *Alb-Cre; Dbp^KI/+^* liver reporter mouse before and after a 6-h advance of the skeleton photoperiod consisting of four 1-h periods of light per 24-h day, as indicated by white. The skeleton photoperiod was advanced by 6 h by shortening the dark phase after the last light pulse on Day 2. Red squares represent the peak of the bioluminescence rhythm, while black circles represent the midpoint of locomotor activity each day, determined by discrete wavelet transform analysis. Six hours of each cycle are double-plotted to aid visualization. Light and dark are indicated by white and gray backgrounds, respectively. **B.** Mean (± SEM) midpoint of locomotor activity (black) and peak of liver bioluminescence (red) rhythms are shown, relative to their initial value, in a group of 4 mice exposed to a 6-h phase advance of the skeleton photoperiod. The locomotor activity rhythm re-sets more rapidly than the bioluminescence rhythm within animal (Significant Measure * Day interaction, and significant phase difference between the rhythms on Day 9; Tukey HSD, *p* <0 .05). **C.** Mean (± SEM) time of midpoint of locomotor activity (black) and peak liver bioluminescence (red) rhythms are shown, relative to their initial phase, in a group of 4 mice not subjected to a phase shift of the skeleton photoperiod.

### Recovery from Circadian Misalignment Induced by Temporally Restricted Feeding

We next conducted a study to examine misalignment induced by restricted feeding. Previous studies have shown that food availability limited to daytime significantly alters phase of peripheral oscillators (Damiola et al., 2000; Hara et al., 2001; Stokkan et al., 2001; Saini et al., 2013). Due to our desire to study bioluminescence rhythms without interference from the LD cycle, we administered different feeding regimens in an LD cycle and then assessed the hepatic bioluminescence rhythm after release to DD with *ad libitum* food. This allowed us to determine the time of peak bioluminescence of the liver after restricted feeding, and the opportunity to continuously observe its return toward a normal phase relationship with SCN-driven behavioral rhythms over time.

*Alb-Cre;Dbp^KI/+^* liver reporter mice were exposed to one of three feeding regimes (*ad libitum*, nighttime, or daytime food availability; **Fig. 7A**) for ten days in LD before recording bioluminescence in DD with *ad libitum* food availability. A previously described automated feeder system (Acosta-Rodriguez et al., 2017) was used to restrict food availability. This system limits total daily consumption (to prevent hoarding) and restricted food pellet delivery for day-fed mice to 0600-1800 h (ZT0-ZT12), and for night-fed mice to 1800-0600 h ZT12 – ZT24/0). With the setting used, the system restored daily food allotments to *ad libitum* fed and night-fed mice daily at 0000h (ZT18), resulting in unusual temporal profiles of food intake in *ad libitum* and night-fed mice. Nevertheless, *ad libitum* and nighttime food access both resulted in food intake being concentrated in the night, while daytime food availability resulted in the midpoint of food intake occurring during the first half of the light phase (**Fig. 7A, 7B, 7C**). Within-group variability in the timing of food intake was low except for three clear outliers (**Fig. 7C**) that were excluded from subsequent analyses.

**Figure 7.**
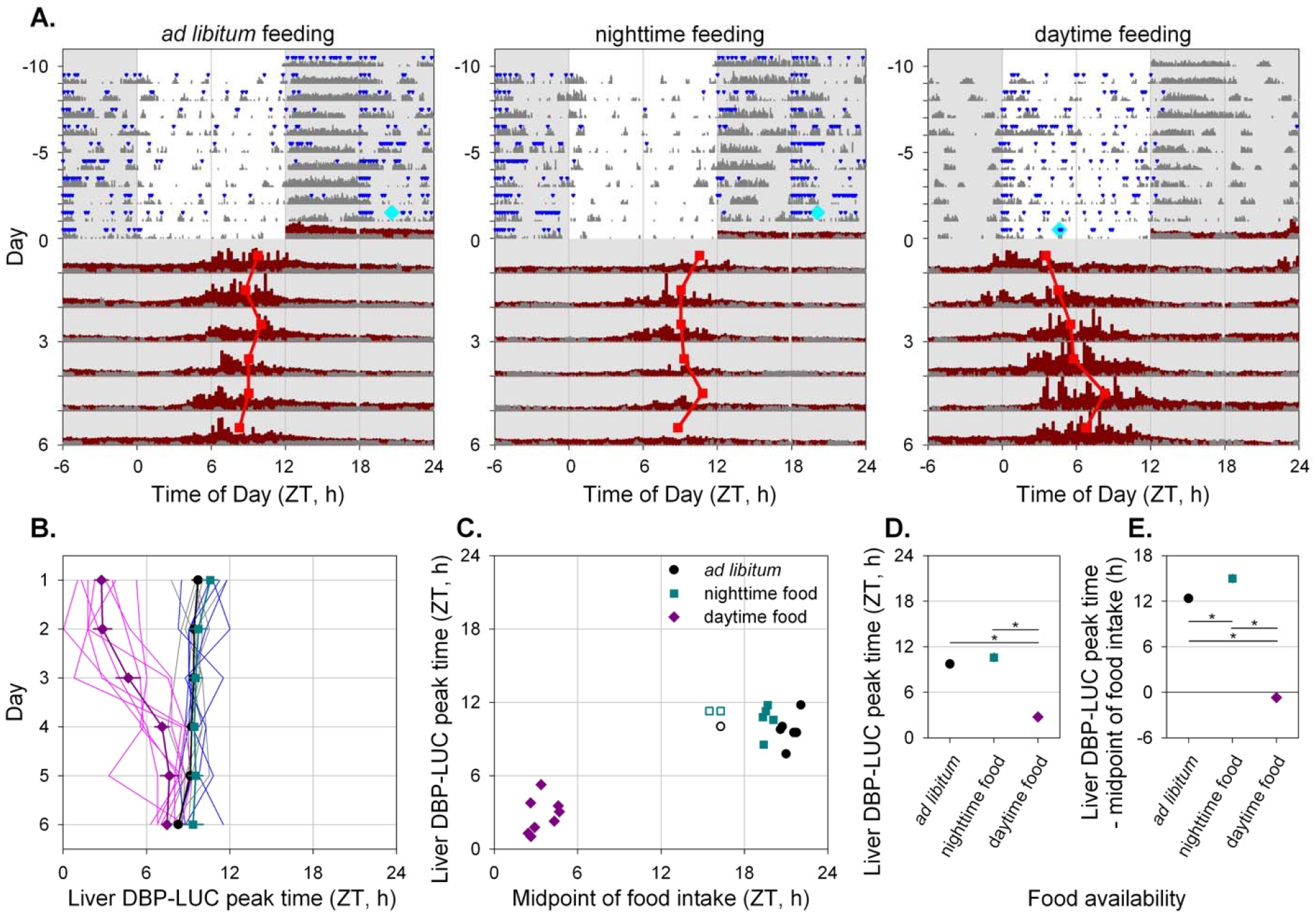
Time-restricted feeding alters the timing of liver bioluminescence rhythms. **A.** Representative actograms of three *Alb-Cre; Dbp^KI/+^* liver reporter mice exposed to the different feeding regimes as indicated above each panel. Mice were housed in 12L:12D lighting and exposed to the specified feeding regime for ten days (-10 to 0) before bioluminescence recording. Food intake (blue triangles) and general locomotor activity (dark gray) were recorded continuously. The midpoint of food intake from days -5 to 0 is indicated by a cyan diamond on day 0. Mice were transferred to the bioluminescence recording setup at the start of the dark phase and housed in constant darkness with *ad libitum* food access. Liver bioluminescence levels are depicted in dark red. Red squares represent the time of peak of the bioluminescence rhythm, determined by DWT. Six hours of each cycle are double-plotted and the y-axis has been stretched during the last 6 days to aid visualization. Light and dark are indicated by white and gray backgrounds, respectively. **B.** Individual and mean (± SEM) phase of liver bioluminescence rhythms relative to clock time for three feeding groups. Mice previously exposed to *ad libitum*, nighttime and daytime feeding are plotted in grey/black, blue/cyan and magenta, respectively (key in Panel C). Prior to recording bioluminescence, mice were entrained to a 12L:12D lighting cycle with lights on at 0600. Mice previously exposed to daytime feeding show an advanced peak phase of liver bioluminescence that reverts over time in constant darkness with *ad libitum* food. **C.** Relationship between preceding feeding phase and peak liver bioluminescence phase for individual animals on the first day under constant conditions. *Ad libitum* and night-fed groups had similar midpoint of food intake; three “outliers” with respect to midpoint of food intake (shown by open symbols) were not included in further analyses (Panels B, D and E). **D.** Mean (± SEM) peak liver bioluminescence phase on the first day under constant conditions, relative to clock time for the three feeding regimens. Error bars were nearly or completely contained within the symbols. **E.** Mean (±SEM) peak liver bioluminescence phase on the first day under constant conditions, relative to the midpoint of preceding food intake for the three feeding regimens. Error bars were nearly or completely contained within the symbols.

*Ad libitum* fed mice showed consistently phased rhythms in bioluminescence after transfer to DD from LD, as did night-fed animals (**Fig. 7A, 7D**). In contrast, mice fed only during the light period for 10 days prior to housing in DD with *ad libitum* food had an earlier peak time of the hepatic bioluminescence rhythm. Daytime feeding resulted in a significantly advanced peak time compared to both night-fed and *ad libitum* fed mice, while these latter groups were statistically indistinguishable (F_2,259.6_ = 76.66, *p* < 0.0001; **Fig. 7D**). Subsequent exposure to DD with *ad libitum* feeding allowed the hepatic clock of day-fed mice to return toward the appropriate phase relationship with the locomotor activity rhythm.

Although daytime feeding resulted in an advanced time of peak bioluminescence, the timing of the liver bioluminescence rhythm was not solely controlled by the timing of food intake. First, no significant correlations between the timing of food intake and time of peak bioluminescence were observed within any of the three feeding regimes (F < 1.13, *p* > 0.32; **Fig. 7C**). Second, the relationship between the timing of liver bioluminescence rhythms relative to the midpoint of food intake was significantly different between the different groups (F_2,17_ = 313.2, p < 0.0001; **Fig. 7E**). While *Dbp*-driven hepatic bioluminescence rhythms were roughly in anti-phase with the midpoint of feeding in *ad libitum* and night-fed mice, daytime feeding resulted in near synchrony between these rhythms (**Fig. 7E**). Furthermore, although the average midpoint of feeding was significantly earlier in night-fed compared to *ad libitum* fed mice (t_10_ = 6.21, p < 0.0001; **Fig. 7C**), no significant difference was observed in bioluminescence phase relative to the preceding light-dark cycle (**Fig. 7D**), with the timing of liver bioluminescence rhythms relative to the midpoint of food intake being significantly delayed in night-fed compared to *ad libitum* fed mice (**Fig. 7E**). Overall, these results demonstrate that although the timing of food intake strongly influences liver rhythms, the timing of bioluminescence rhythmicity in liver reporter mice is not solely driven by the timing of food intake (with food intake regulated for this duration and in this way).

## Discussion

Numerous studies have made use of rhythmically expressed bioluminescent reporter genes to monitor circadian rhythms. The *Per2^Luiferase^* mouse and other reporters with bioluminescence under the control of a clock gene have been especially useful as they generate robust bioluminescence rhythms from numerous tissues recorded *ex vivo* (Abe et al., 2002; Maywood et al., 2013; Yakazami et al., 2000; Yamazaki and Takahashi, 2005; Yoo et al 2004; Yoo et al., 2005). The widespread expression of PER2::LUCIFERASE (and other ‘non-conditional’ bioluminescence reporters) comes at a cost, however, as it is not possible to assess rhythmicity in specific cell populations within a larger tissue without dissection. Tissue explant preparation can cause phase-resetting, however, especially after exposure to phase shifting stimuli (Noguchi et al., 2020; Leise et al., 2020). Furthermore, *ex vivo* culturing of tissues does not allow assessment of rhythmicity in the context of the hierarchical circadian system or dynamic changes during environmentally-induced resetting.

Addressing issues of internal desynchrony and misalignment of oscillators requires monitoring the dynamics of tissue resetting over time after a phase-shifting stimulus. The use of *in vivo* bioluminescence imaging for repeated assessments of organ-level regions of interest over multiple days is feasible but requires several potentially disruptive anesthesia sessions (Poulsen et al., 2018) per circadian cycle. As a result, *in vivo* bioluminescence imaging has generally been relegated to assessing phase of reporter gene oscillations on relatively few occasions after a shifting stimulus, with rare exception (van der Vinne et al., 2020). Other methods for monitoring bioluminescence and fluorescence rhythms in ambulatory mice have been developed (Hamada et al., 2016; Mei et al., 2018; Nakamura et al., 2008; Ono et al., 2015; Saini et al., 2013; Sawai et al., 2019; Yamaguchi et al., 2016; Yamaguchi et al., 2001), but a less invasive approach for assessing rhythms in a variety of specific tissues is desirable. Notably, several abdominal organs emit significant amounts of bioluminescence in “whole-body” reporter mice, including liver, kidney and intestines. These tissues likely overshadow (or, more accurately, out-glow) surrounding tissues. Bioluminescence from even larger organs like liver and kidney is likely ‘contaminated’ by light from adjacent structures. Indeed, the size and shape of the “liver” ROI seen by IVIS imaging (Fig. 4) differs between *Dbp* liver reporter mice and whole-body reporter *Dbp^Luc^* mice. These considerations underline the benefits of generating a *Cre*-conditional reporter mouse in which recombination leading to bioluminescence can be directed to specific tissues and cell types.

We chose to modify the *Dbp* gene to generate a conditional reporter for several reasons. *Dbp* is widely and rhythmically expressed at readily detectable levels (Fonjallaz et al., 1996; Punia et al., 2012; Zhang et al., 2014). This feature ensures that the reporter mouse would be useful for detecting rhythmicity in numerous tissues. In addition, individual clock genes are responsive to different signaling pathways. This differential regulation can lead to circadian misalignment *within* the circadian clock (Reddy et al., 2002; Nicholls et al., 2019). As an output gene controlled by the CLOCK:BMAL1 transcriptional activator complex (Ripperger & Schibler 2006; Stratmann et al., 2012), *Dbp* rhythmicity is likely a good proxy for the integrated output of the molecular clockwork. Additional *cis*-acting elements regulating *Dbp* expression have been identified, however. Binding of hnRNP K to a poly-(C) motif in the proximal promoter has been implicated in high-amplitude expression of *Dbp* (Kwon et a., 2019; Kwon et al., 2020). Interestingly, *Dbp* appears to be insensitive to acute regulation by activation of signal transduction pathways. Unlike *Per1* and *Per2*, *Dbp* gene expression is not increased in the mouse SCN following exposure to light at night (Yan et al., 2000). Furthermore, *Dbp* expression is not acutely increased by horse serum or stimulation of the cAMP/PKA pathway by forskolin, which rapidly induce *Per1* (Yagita & Okamura, 1999) and resynchronize molecular rhythms. Thus, *Dbp* expression and the *Dbp*-based reporter are likely to represent the status of the molecular clock without interference by other influences. Finally, concern that the targeting event could disrupt function of the modified gene led us to steer away from core clock genes. Mice homozygous for a targeted allele of *Dbp* have only a modest circadian phenotype (Lopez-Molina et al., 1997). Homozygotes of both the *Per2^LucSV^* and *Per2^Luciferase^* lines have altered circadian rhythms (Ralph et al., 2020; Yoo et al., 2017; see below). The GFP-expressing *Dbp* transcript lacks the native 3′ UTR and uses an exogenous polyadenylation sequence, which could affect *Dbp* gene expression and regulation. Notably, however, our Northern blot analysis suggests little or no alteration in expression level or dynamics of the *Dbp* reporter transcripts.

Yoo et al. (2017) reported that homozygous *Per2^LucSV/LucSV^* mice (in which a SV40 polyadenylation site is used instead of the endogenous *Per2* 3′ UTR) have a longer period length of locomotor activity rhythms in DD and explant bioluminescence rhythms *ex vivo* than the more widely used *Per2^Luciferase^* reporter. The potential impact of a single *Per2^LucSV^* allele (as used in our studies) on period length has not been reported, but this could contribute to the longer period of *Per2^LucSV^*^/+^ explants, relative to *Dbp^Luc/+^* explants. Interestingly, Ralph et al., (2020) recently reported that the *Per2^Luciferase^* reporter that uses the endogenous *Per2* 3′UTR (Yoo et al., 2004) also has longer period in DD and other circadian phenotypes. Tosini and colleagues have also recently reported retinal degeneration and alterations in classical photoreception in aged male *Per2^Luciferase/Liuciferase^* mice (Goyal et al., 2021).

Shan et al. (2020) recently reported development of a Color-Switch PER2::LUC line which was used to demonstrate the utility of a *Cre*-dependent reporter approach for interrogating SCN circuitry. The Color-Switch PER2::Luc line has the advantage of reporting on both *Cre*-positive and *Cre*-negative cells in different colors. Detection of bioluminescence from the Color-Switch PER2::LUC reporter requires segmentation of the bioluminescence signal between wavelengths. Our ‘simpler’ approach of only inducing a bioluminescence signal in *Cre*-positive cells of *Dbp^KI/+^* mice enables recording of bioluminescence rhythms without the need for wavelength segmentation. In addition, the *Dbp* reporter can easily be used in *Per2* mutant mice. Like the Color-Switch PER2::LUC line, our *Dbp* conditional reporter line is useful for *ex vivo* studies, allowing specific cellular populations to be monitored by crossing to the appropriate *Cre-*expressing lines.

As with the Color-Switch PER2::LUC line, we also intended to generate a bifunctional reporter. The inability to readily detect a GFP fluorescence rhythm in *Dbp^KI/+^* mouse SCN or liver was unexpected. It is important to emphasize that there was detectable fluorescence above baseline, but rhythmicity was not detected. This could nevertheless be due in part to low signal-to-noise ratio. It is possible that the short half-life of destabilized GFP, combined with the waveform of *Dbp* gene expression (being less sinusoidal than the rhythms of *Per1* and *Per2*, for example) contributed to a short period of production and rapid degradation of the GFP. Notably, a transgenic mouse in which a similarly destabilized GFP is driven by the *Per1* promoter generates nice SCN fluorescence rhythms (Kuhlman et al., 2000). Similarly, destabilized versions of VENUS (a yellow fluorescent protein) and DsRED inserted at the start codon of PER1 and PER2, respectively, in bacterial artificial chromosomes have been useful for monitoring rhythmic gene expression in SCN of transgenic mice (Cheng et al., 2009). Assessing immunoreactive GFP (rather than native fluorescence from GFP) in the *Dbp^KI/+^* or *Dbp^KI/KI^* mice may improve signal detection (but at the cost of real-time reporting).

It is possible that a different design of the reporter construct would have led to better success. Addition of a nuclear localization signal (as done by Cheng et al., 2009) would reduce the volume in which GFP is distributed, making signal intensity greater (but we note that Kuhlman et al., 2000 did not incorporate an NLS into their reporter sequence). Alternatively, generating non-destabilized GFP as a fusion protein with DBP might have been more successful; in this scenario, the stability of DBP would regulate the stability of GFP. This fusion strategy has been used successfully with fluorescent reporters of PER2 and BMAL1 (Smyllie et al., 2016; Yang et al., 2020). Another potential variation is to use fluorescent proteins other than GFP. Other fluorescent proteins may be brighter and thus more amenable to this type of study.

Our studies reveal subtle differences among the population of oscillators defined by AVP-*Cre*, NMS-*Cre*, and the entire SCN cohort. More specifically, AVP cells had a shorter period, reduced rhythmicity index, and larger within-slice dispersal of peak times than the NMS cell population with which it overlaps. Our results suggest that AVP cells are coordinated less well and are less robust than other populations in the SCN. This suggestion is in contrast to the typical view of AVP cells as high-amplitude ‘output’ neurons that also contribute to determination of period and rhythm amplitude (Herzog et al., 2017; Mieda et al., 2015; Mieda et al., 2016). One possible explanation for this is that AVP is dysregulated in the *Avp-Cre* line (Cheng et al., 2019), which may influence the function of the SCN as a whole in the *AVP-IRES-Cre; Dbp^KI/+^* genotype used here. Using an *Avp-IRES-Cre* line which does not reduce AVP expression, Shan et al. (2020) reported that AVP-expressing SCN neurons have shorter period bioluminescence rhythms, compared to the non-AVP cells. This contrasts directly with our finding of longer period in AVP cells reporting luciferase from the *Dbp* locus. The AVP neuronal population is contained entirely within the NMS-expressing population in the SCN. There is no evidence that the *Nms-Cre* line alters circadian timekeeping on its own (Lee et al., 2015). The *Nms-Cre* line and the *Avp-IRES-Cre* line used by Shan et al. (2020) appear to be preferable models to the AVP-IRES-*Cre* line (JAX 023530) used here. Of note for circadian researchers, a *Vip-IRES-Cre* line also influences neuropeptide expression and circadian function (Cheng et al., 2019; Joye et al., 2020).

We envision this line will be very useful for monitoring additional neuronal subpopulations in the SCN in wild-type and mutant animals. Additional technical development may allow *in vivo* detection of bioluminescence rhythms from neuronal populations in awake behaving mice. Approaches to optimize the signal detected from brain include use of highly efficient and cell- and brain-penetrant substrates (Evans et al., 2014; Iwano et al., 2018), cranial windows (Miller et al., 2014) and hairless or albino mice (Martin-Burgos et al., 2020; Iwano et al., 2018). (Note, the tyrosinase mutation leading to albinism in C57BL/6J mice is linked to *Dbp* on mouse chromosome 7; we have nevertheless generated recombinants and produced albino reporters, including the kidney reporter mice in **Fig. S3**). These approaches may allow interrogation of the SCN circuit *in vivo*, extending the elegant studies being performed with SCN slices *ex vivo.* Bioluminescence rhythms can also be examined in neuronal populations outside the SCN, by using an appropriate *Cre* driver and/or viral delivery of *Cre* recombinase.

*Cre*-mediated recombination of the *Dbp^KI^* allele in liver enabled us to perform continuous, *in vivo* bioluminescence monitoring of liver in freely moving mice. These studies demonstrate transient misalignment between the liver oscillator and SCN-regulated behavioral rhythms. Our design is complementary to that used by Saini et al. (2013), who continuously monitored reporter gene bioluminescence as hepatic rhythms were shifted by an inverted feeding regimen.

Repeated misalignment among oscillators is thought to contribute to adverse metabolic and health consequences of chronic circadian disruption (for reviews, see Arble et al., 2015; Evans and Davidson, 2013; Roenneberg and Merrow, 2016; Patke et al., 2020; West & Bechtold, 2015). Up until now, technical and practical limitations have restricted our ability to monitor the behavior of circadian rhythms in different peripheral tissues during and following environmental disruption of circadian homeostasis. Our *Cre*-conditional reporter line and the approaches described recently (Martin-Burgos et al., 2022; Tam et al., 2021), and extended here for longitudinal and tissue-specific assessment of bioluminescence rhythms *in vivo* will allow characterization of misalignment and recovery after a variety of circadian-disruptive lighting and food availability paradigms. These approaches will allow more extensive examination of the consequences of repeated misalignment of peripheral clocks.

The data in Figure 6 show clear misalignment between the rhythms in locomotor activity and hepatic bioluminescence. Rather remarkably, the average phase of peak bioluminescence did not shift at all for 3-4 days after the shift of the lighting cycle. This is consistent with previous work by others: even with a more robust signal that directly impacts peripheral oscillators (reversal of the time of food availability), shifting of the liver clock occurs slowly (Damiola et al., 2001; Saini et al., 2013). We did not track food intake in this experiment, so do not know the rate at which the food intake pattern was reset following the shift of the lighting cycle. The timing of food intake is typically controlled by the SCN, however, and thus would likely track locomotor activity. The food intake and locomotor activity rhythms shift to the new phase more rapidly than the hepatic oscillator, resulting in misalignment.

Previous studies have shown misalignment between central and peripheral clocks induced by altering the time of food access to daytime, by assessing oscillator phase at various time-points after a phase shift of the lighting cycle, or by exposure to non-24hr light-dark schedules. The vast majority of these studies monitored bioluminescence rhythms *ex vivo* or assessed transcript levels following tissue collection at various times after a shift (Balsalobre et al., 2000; Damiola et al., 2000; Davidson et al., 2008; Davidson et al., 2009; Nakamura et al., 2005; Nicholls et al., 2019; Pezuk et al., 2012; Sellix et al., 2012; Stokkan et al., 2001; Yamanaka et al., 2008). Notably, *ex vivo* bioluminescence rhythm timing may be affected by prior lighting conditions (Noguchi et al., 2020; Leise et al., 2020; Tahara et al., 2012). Few studies have followed bioluminescence rhythms *in vivo* over time after a light-induced phase shift or after a food manipulation that phase-shifts peripheral oscillators (but see Saini et al., 2013; van der Vinne et al., 2020). Our current data leverage the ability to non-invasively monitor rhythmicity from a single peripheral oscillator in individual animals over many days to show the time course of internal misalignment and recovery after a phase shift. Other studies with minimally invasive monitoring of bioluminescence rhythms have relied upon viral introduction of the reporter into liver, and thus cannot easily be extended to other tissues (Saini et al., 2013; Sinturel et al., 2021). Notably, the viral reporter appears not strictly limited to liver in this model (see Saini et al. 2013, their Fig S2). Moreover, efficient expression of virally delivered reporter constructs is limited by the promoter size and specificity, so the level and anatomical pattern of expression often do not match that of the gene whose promoter was used. Future studies of additional tissues in *Cre*-conditional reporter mice will enable elucidation of how other tissues within the hierarchical, multi-oscillatory circadian system respond to disruptive stimuli. Several studies suggest that organs differ in their response to resetting stimuli. For example, the *Dbp* mRNA rhythm in liver is more fully reset than the rhythm in heart and kidneys 3 days after restricting food availability to daytime (Damiola et al., 2000), and several studies indicate the SCN (and the locomotor rhythms it regulates) resets more rapidly than peripheral tissues (Davidson et al., 2008; Davidson et al., 2009; Damiola et al., 2000; Hamada et al., 2016; Saini et al., 2013; Sellix et al., 2012; van der Vinne et al., 2020; Yamanaka et al., 2008; Yamazaki et al., 2000; see Nicholls et al., 2019 for review).

A recent study used a feeding device similar to the one used here (Acosta-Rodriquez et al, 2017) to recapitulate ‘naturalistic’ food intake patterns in mice with restricted food access (Xie et al., 2020). In this study, the food restriction was not the severe ‘all or none’ patterns typically used in studies with time-restricted access to food. The authors found that peripheral oscillators of *Per2^Luciferase^* mice were not effectively entrained by restricted feeding using the imposed ‘natural’ feeding patterns (Xie et al., 2020). Similarly, our study (shown in Figure 7) revealed that daytime restricted food access produced a smaller and more variable phase shift of the hepatic circadian clock (as indicated by the initial time-of-peak of *Dbp*-driven bioluminescence) than expected based on published results using presence / absence food availability cycles (Damiola et al., 2000; Hara et al., 2001; Saini et al., 2013; Stokkan et al., 2001). Both day-to-day variation in phase of peak bioluminescence within animals as well as variation in peak phase between animals is larger in the day-fed mice than in the night-fed and *ad lib* groups. These latter two groups did not need to change the time of food intake greatly, while the daytime-fed group was eating at an abnormal phase. Our imposing temporal restriction on feeding for only 10 days before release to DD and *ad lib* feeding may not have been sufficient to synchronize liver clocks to a new phase, as suggested by the partial reversal of the phase of hepatic bioluminescence. In addition, food intake patterns derived from presence/absence cycles of food appear much more effective at synchronizing the liver than more naturalistic food intake patterns (our data and Xie et al., 2020 compared with, for example, Saini et al., 2013 and Damiola et al., 2001). The night-fed and *ad lib* groups have relatively more intense, consolidated feeding in the early part of the night, which may provide a stronger stimulus to peripheral oscillators, including liver. Food access for 12 hours during the daytime may be less concentrated and more variable in time, providing a less effective synchronizing cue to peripheral oscillators. This may lead to higher levels of within-organ desynchrony among cells and thus lower-amplitude rhythmicity, secondarily leading to greater variability in determining the time of peak bioluminescence on subsequent days. We also cannot rule out the possibility that the time of food intake differed between individuals and between the groups, even during *ad lib* feeding, and this could influence hepatic rhythms. Future studies using a shorter duration of food access per day and monitoring bioluminescence rhythms both during the acclimation to daytime feeding as well as during release to *ad lib* conditions, coupled with monitoring food intake patterns throughout the study, should allow more dynamic assessment of the entrainment and subsequent free-running rhythms of peripheral oscillators *in vivo.* In addition, use of a variety of different *Cre* drivers will allow assessment of whether different peripheral organs respond similarly to food restriction paradigms. In addition, tissue-specific reporter models will be very useful in assessing how more naturalistic food ingestion paradigms influence peripheral circadian clocks in several tissues.

In summary, we have demonstrated the utility of a new, *Cre*-conditional reporter mouse that enables tissue-specific monitoring of circadian molecular rhythms *in vivo* and *ex vivo*. This reporter mouse provides a major advance in our capabilities for monitoring rhythms in a variety of tissues under normal and disruptive conditions, which is a key step in the identification of mechanisms underlying the adverse consequences of circadian disruption inherent to life in modern 24/7 societies.

## Acknowledgments

We thank Christopher Lambert and Jamie Black for technical assistance, and Steven A. Brown (University of Zurich) for discussions during the development of this project. UMass Chan Medical School core facilities (Mutagenesis Core, Mouse Modeling Core, and Small Animal Imaging Core) are gratefully acknowledged.

Research reported in this publication was supported by the National Institute for Neurological Diseases and Stroke and the National Institute of General Medical Sciences of the National Institutes of Health under award numbers R21NS103180 (DRW), SC1GM112567 (AJD), and NIGMS R15GM126545 (MEH), the Hartmann Müller Stiftung (RD), MRC MC_PC_15070 (RD) and BSN (RD and LAG). CBS was a participant in the UMass Chan Medical School Initiative for Maximizing Student Development, supported by NIH grant R25GM113686. The funders had no role in study design, data collection and analysis, decision to publish, or preparation of the manuscript. The content is solely the responsibility of the authors and does not necessarily represent the official views of the National Institutes of Health or the other funding agencies.

## Author Contributions

R.D and D.R.W. conceived the project

C.B.S., V.v.d.V., E.M., M.H.B., A.J.D., M.E.H., R.D. and D.R.W. designed research

C.B.S., V.v.d.V., E.M., A.C.S., B.M.B., P.C.M., L.A.G., R.D., and D.R.W. performed research

C.B.S., V.v.d.V., E.M., T.L.L., B.M.B., M.E.H., R.D. and D.R.W. analyzed data

C.B.S., V.v.d.V., and D.R.W. wrote the paper

All authors have approved this version of the manuscript.

## Notes

**Conflict of interest statement:** The authors declare no conflicts of interest.

### Competing Interest Statement

The authors have declared no competing interest.

### Summary of Updates

Incorporates response to reviewer concerns and comments, in Discussion section. Addition of Supplemental figure S7.

## References

Abe M, Herzog ED, Yamazaki S, Straume M, Tei H, Sakaki Y, Menaker M, and Block GD (2002) Circadian rhythms in isolated brain regions. J Neurosci 22:350–356.

Acosta-Rodriguez VA, de Groot, M H M, Rijo-Ferreira F, Green CB, and Takahashi JS (2017) Mice under caloric restriction self-impose a temporal restriction of food intake as revealed by an automated feeder system. Cell Metab 26:267–277.e2.

Balsalobre A, Brown SA, Marcacci L, Tronche F, Kellendonk C, Reichardt HM, Schutz G, and Schibler U (2000) Resetting of circadian time in peripheral tissues by glucocorticoid signaling. Science 289:2344–2347.

Brandes C, Plautz JD, Stanewsky R, Jamison CF, Straume M, Wood KV, Kay SA, and Hall JC (1996) Novel features of *Drosophila period* transcription revealed by real-time luciferase reporting. Neuron 16:687–692.

Cesbron F, Brunner M, and Diernfellner AC (2013) Light-dependent and circadian transcription dynamics *in vivo* recorded with a destabilized luciferase reporter in *Neurospora*. PLoS One 8:e83660.

Chen Z, Yoo SH, Park YS, Kim KH, Wei S, Buhr E, Ye ZY, Pan HL, and Takahashi JS (2012) Identification of diverse modulators of central and peripheral circadian clocks by high-throughput chemical screening. Proc Natl Acad Sci U S A 109:101–106.

Cheng AH, Fung SW, and Cheng HM (2019) Limitations of the AVP-IRES2-Cre (JAX #023530) and VIP-IRES-Cre (JAX #010908) models for chronobiological investigations. J Biol Rhythms 34:634–644.

Damiola F, Le Minh N, Preitner N, Kornmann B, Fleury-Olela F, and Schibler U (2000) Restricted feeding uncouples circadian oscillators in peripheral tissues from the central pacemaker in the suprachiasmatic nucleus. Genes Dev 14:2950–2961.

Davidson AJ, Castanon-Cervantes O, Leise TL, Molyneux PC, and Harrington ME (2009) Visualizing jet lag in the mouse suprachiasmatic nucleus and peripheral circadian timing system. Eur J Neurosci 29:171–180.

Davidson AJ, Yamazaki S, Arble DM, Menaker M, and Block GD (2008) Resetting of central and peripheral circadian oscillators in aged rats. Neurobiol Aging 29:471–477.

Destici E, Jacobs EH, Tamanini F, Loos M, van der Horst GTJ, Oklejewicz M (2013) Altered phase-relationship between peripheral oscillators and environmental time in *Cry1* or *Cry2* deficient mouse models for early and late chronotypes. PLoS ONE 8, e83802.

Evans JA and Davidson AJ (2013) Health consequences of circadian disruption in humans and animal models. Prog Mol Biol Transl Sci 119:283–323.

Evans JA, Leise TL, Castanon-Cervantes O, and Davidson AJ (2013) Dynamic interactions mediated by nonredundant signaling mechanisms couple circadian clock neurons. Neuron 80:973–983.

Evans JA, Leise TL, Castanon-Cervantes O, and Davidson AJ (2011) Intrinsic regulation of spatiotemporal organization within the suprachiasmatic nucleus. PLoS One 6:e15869.

Evans MS, Chaurette JP, Adams ST, Reddy GR, Paley MA, Aronin N, Prescher JA, and Miller SC (2014) A synthetic luciferin improves bioluminescence imaging in live mice. Nat Methods 11:393–395.

Fonjallaz P, Ossipow V, Wanner G, and Schibler U (1996) The two PAR leucine zipper proteins, TEF and DBP, display similar circadian and tissue-specific expression, but have different target promoter preferences. EMBO J 15:351–362.

Goyal V, DeVera C, Baba K, Sellers J, Chrenek MA, Iuvone PM, Tosini G (2021) Photoreceptor degeneration in homozygous male *Per2^luc^* mice during aging. J Biol Rhythms 36:137–145.

Hamada T, Sutherland K, Ishikawa M, Miyamoto N, Honma S, Shirato H, and Honma K (2016) *In vivo* imaging of clock gene expression in multiple tissues of freely moving mice. Nat Commun 7:11705.

Hara R, Wan K, Wakamatsu H, Aida R, Moriya T, Akiyama M, and Shibata S (2001) Restricted feeding entrains liver clock without participation of the suprachiasmatic nucleus. Genes Cells 6:269–278.

Harris JA, Hirokawa KE, Sorensen SA, Gu H, Mills M, Ng LL, Bohn P, Mortrud M, Ouellette B, Kidney J, Smith KA, Dang C, Sunkin S, Bernard A, Oh SW, Madisen L, and Zeng H (2014) Anatomical characterization of Cre driver mice for neural circuit mapping and manipulation. Front Neural Circuits 8:76.

Herzog ED hermanstyne T, Smyllie NJ, Hastings MH (2017) Regulating the suprachiasmatic nucleus (SCN) clockwork: Interplay between cell-autonomous and circuit-level mechanisms. Cold Springs Harbor Perspect Biol 9: a027706.

Hirota T, Lee JW, Lewis WG, Zhang EE, Breton G, Liu X, Garcia M, Peters EC, Etchegaray JP, Traver D, Schulz PG, Kay SA (2010) High-throughput chemical screen identified a novel potent modulator of cellular circadian rhythms and reveals CK1 alpha as a clock regulatory kinase. PLoS Biol 8, e1000559

Iwano S, Sugiyama M, Hama H, Watakabe A, Hasegawa N, Kuchimaru T, Tanaka KZ, Takahashi M, Ishida Y, Hata J, Shimozono S, Namiki K, Fukano T, Kiyama M, Okano H, Kizaka-Kondoh S, McHugh TJ, Yamamori T, Hioki H, Maki S, and Miyawaki A (2018) Single-cell bioluminescence imaging of deep tissue in freely moving animals. Science 359:935–939.

Joye DAM, Rohr KE, Keller D, Inda T, Telega A, Pancholi H, Carmona-Alcocer V, and Evans JA (2020) Reduced VIP expression affects circadian clock function in VIP-IRES-CRE Mice (JAX 010908). J Biol Rhythms 35:340–352.

Kim JH, Lee SR, Li LH, Park HJ, Park JH, Lee KY, Kim MK, Shin BA, Choi SY. (2011) High cleavage efficiency of a 2A peptide derived from porcine Teschovirus-1 in human cell lines, zebrafish and mice. PLoS ONE 6, e18556

Kondo T, Strayer CA, Kulkarni RD, Taylor W, Ishiura M, Golden SS, and Johnson CH (1993) Circadian rhythms in prokaryotes: luciferase as a reporter of circadian gene expression in cyanobacteria. Proc Natl Acad Sci U S A 90:5672–5676.

Kuhlman SJ, Quintero JE, McMahon DG (2000) GFP fluorescence reports *Period 1* circadian gene regulation in the mammalian biological clock. NeuroReport 11:1479–1482.

Kwon PK, Kim H-M, Kim SW, Kang B, Yi H, Ku H-O, Roh T-Y, Kim K-T (2019) The poly(C) motif in the proximal promoter region of D site-binding protein gene (Dbp) drives its high-amplitude oscillation. Mol Cell Biol 39:e00101–19

Kwon PK, Lee K-H, Kim J-h, Tae S, Ham S, Jeong Y-H, Kim SW, Kang B, Kim H-M, Choi J-H, Yi H, Ku H-O, Roh T-Y, Lim C, Kim K-T (2020) hnRNP K supports high-amplitude D site-binding protein mRNA (*Dbp* mRNA) oscillation to sustain circadian rhythms. Mol Cell Biol 40:e00537–19.

Lee IT, Chang AS, Manandhar M, Shan Y, Fan J, Izumo M, Ikeda Y, Motoike T, Dixon S, Seinfeld JE, Takahashi JS, and Yanagisawa M (2015) Neuromedin S-producing neurons act as essential pacemakers in the suprachiasmatic nucleus to couple clock neurons and dictate circadian rhythms. Neuron 85:1086–1102.

Leise TL (2017) Analysis of nonstationary time series for biological rhythms research. J Biol Rhythms 32:187–194.

Leise TL, Goldberg A, Michael J, Montoya G, Solow S, Molyneux P, Vetrivelan R, and Harrington ME (2020) Recurring circadian disruption alters circadian clock sensitivity to resetting. Eur J Neurosci 51:2343–2354.

Leise TL and Harrington ME (2011) Wavelet-based time series analysis of circadian rhythms. J Biol Rhythms 26:454–463.

Leise TL, Harrington ME, Molyneux PC, Song I, Queenan H, Zimmerman E, Lall GS, and Biello SM (2013) Voluntary exercise can strengthen the circadian system in aged mice. Age (Dordr) 35:2137–2152.

Logan M, Martin JF, Nagy A, Lobe C, Olson EN, and Tabin CJ (2002) Expression of Cre recombinase in the developing mouse limb bud driven by a Prxl enhancer. Genesis 33:77–80.

Lopez-Molina L, Conquet F, Dubois-Dauphin M, and Schibler U (1997) The DBP gene is expressed according to a circadian rhythm in the suprachiasmatic nucleus and influences circadian behavior. EMBO J 16:6762–6771.

Martin-Burgos B, Wang W, William I, Tir S, Mohammad I, Javed R, Smith S, Cui Y, Smith CB, van der Vinne V, Molyneux PC, Miller SC, Weaver DR, Leise TL, Harrington ME (2020) Methods for detecting PER2::LUCIFERASE bioluminescence rhythms in freely moving mice. BioRxiv https://doi.org/10.1038/s41467-017-00462-2. Journal of Biological Rhythms, in press 2022.

Maywood ES, Drynan L, Chesham JE, Edwards MD, Dardente H, Fustin JM, Hazlerigg DG, O’Neill JS, Codner GF, Smyllie NJ, Brancaccio M, and Hastings MH (2013) Analysis of core circadian feedback loop in suprachiasmatic nucleus of *mCry1-luc* transgenic reporter mouse. Proc Natl Acad Sci U S A 110:9547–9552.

Mei L, Fan Y, Lv X, Welsh DK, Zhan C, and Zhang EE (2018) Long-term in vivo recording of circadian rhythms in brains of freely moving mice. Proc Natl Acad Sci U S A 115:4276–4281.

Mieda M, Okamoto H, and Sakurai T (2016) Manipulating the cellular circadian period of arginine vasopressin neurons alters the behavioral circadian period. Curr Biol 26:2535–2542.

Mieda M, Ono D, Hasegawa E, Okamoto H, Honma K, Honma S, and Sakurai T (2015) Cellular clocks in AVP neurons of the SCN are critical for interneuronal coupling regulating circadian behavior rhythm. Neuron 85:1103–1116.

Millar AJ, Short SR, Chua NH, and Kay SA (1992) A novel circadian phenotype based on firefly luciferase expression in transgenic plants. Plant Cell 4:1075–1087.

Millar AJ, Carre IA, Strayer CA, Chua NH, Kay SA (1995). Circadian clock mutants in Arabidopsis identified by luciferase imaging. Science 267:1161–1163.

Miller JE, Granados-Fuentes D, Wang T, Marpegan L, Holy TE, and Herzog ED (2014) Vasoactive intestinal polypeptide mediates circadian rhythms in mammalian olfactory bulb and olfaction. J Neurosci 34:6040–6046.

Mohawk JA, Green CB, and Takahashi JS (2012) Central and peripheral circadian clocks in mammals. Annu Rev Neurosci 35:445–462.

Morgan LW, Greene AV, and Bell-Pedersen D (2003) Circadian and light-induced expression of luciferase in *Neurospora crassa*. Fungal Genet Biol 38:327–332.

Muñoz-Guzmán F, Caballero V, and Larrondo LF (2021) A global search for novel transcription factors impacting the *Neurospora crassa* circadian clock. G3 (Bethesda). DOI: 10.1093/g3journal/jkab100

Nagano M, Adachi A, Nakahama K, Nakamura T, Tamada M, Meyer-Bernstein E, Sehgal A, and Shigeyoshi Y (2003) An abrupt shift in the day/night cycle causes desynchrony in the mammalian circadian center. J Neurosci 23:6141–6151.

Nagoshi E, Saini C, Bauer C, Laroche T, Naef F, and Schibler U (2004) Circadian gene expression in individual fibroblasts: cell-autonomous and self-sustained oscillators pass time to daughter cells. Cell 119:693–705.

Nakamura W, Yamazaki S, Takasu NN, Mishima K, and Block GD (2005) Differential response of *Period 1* expression within the suprachiasmatic nucleus. J Neurosci 25:5481–5487.

Nakamura W, Yamazaki S, Nakamura TJ, Shirakawa T, Block GD, and Takumi T (2008) *In vivo* monitoring of circadian timing in freely moving mice. Curr Biol 18:381–385.

Nicholls SK, Casiraghi LP, Wang W, Weber ET, and Harrington ME (2019) Evidence for internal desynchrony caused by circadian clock resetting. Yale J Biol Med 92:259–270.

Noguchi T, Harrison EM, Sun J, May D, Ng A, Welsh DK, and Gorman MR (2020) Circadian rhythm bifurcation induces flexible phase resetting by reducing circadian amplitude. Eur J Neurosci 51:2329–2342.

Ono D, Honma K, and Honma S (2015a) Circadian and ultradian rhythms of clock gene expression in the suprachiasmatic nucleus of freely moving mice. Sci Rep 5:12310.

Patke A, Young MW, and Axelrod S (2020) Molecular mechanisms and physiological importance of circadian rhythms. Nat Rev Mol Cell Biol 21:67–84.

Pezuk P, Mohawk JA, Wang LA, and Menaker M (2012) Glucocorticoids as entraining signals for peripheral circadian oscillators. Endocrinology 153:4775–4783.

Postic C, Shiota M, Niswender KD, Jetton TL, Chen Y, Moates JM, Shelton KD, Lindner J, Cherrington AD, and Magnuson MA (1999) Dual roles for glucokinase in glucose homeostasis as determined by liver and pancreatic beta cell-specific gene knock-outs using Cre recombinase. J Biol Chem 274:305–315.

Poulsen RC, Warman GR, Sleigh J, Ludin NM, and Cheeseman JF (2018) How does general anaesthesia affect the circadian clock? Sleep Med Rev 37:35–44.

Punia S, Rumery KK, Yu EA, Lambert CM, Notkins AL, and Weaver DR (2012) Disruption of gene expression rhythms in mice lacking secretory vesicle proteins IA-2 and IA-2β. Am J Physiol Endocrinol Metab 303:762.

Ralph MR, Shi SQ, Johnson CH, Houdek P, Shrestha TC, Crosby P, O’Neill JS, Sladek M, Stinchcombe AR, and Sumova A (2021) Targeted modification of the *Per2* clock gene alters circadian function in *mPer2^luciferase^* (*mPer2^Luc^*) mice. PLoS Comput Biol 17:e1008987.

Reddy AB, Field MD, Maywood ES, and Hastings MH (2002) Differential resynchronisation of circadian clock gene expression within the suprachiasmatic nuclei of mice subjected to experimental jet lag. J Neurosci 22:7326–7330.

Ripperger JA and Schibler U (2006) Rhythmic CLOCK-BMAL1 binding to multiple E-box motifs drives circadian Dbp transcription and chromatin transitions. Nat Genet 38:369–374.

Ripperger JA, Shearman LP, Reppert SM, Schibler U (2000) CLOCK, an essential pacemaker component, controls expression of the circadian transcription factor DBP. Genes Dev 14:679–689.

Roenneberg T and Merrow M (2016) The Circadian Clock and Human Health. Curr Biol 26:432.

Saini C, Liani A, Curie T, Gos P, Kreppel F, Emmenegger Y, Bonacina L, Wolf JP, Poget YA, Franken P, and Schibler U (2013) Real-time recording of circadian liver gene expression in freely moving mice reveals the phase-setting behavior of hepatocyte clocks. Genes Dev 27:1526–1536.

Sawai Y, Okamoto T, Muranaka Y, Nakamura R, Matsumura R, Node K, and Akashi M (2019) *In vivo* evaluation of the effect of lithium on peripheral circadian clocks by real-time monitoring of clock gene expression in near-freely moving mice. Sci Rep 9:10909–3.

Sellix MT, Evans JA, Leise TL, Castanon-Cervantes O, Hill DD, DeLisser P, Block GD, Menaker M, and Davidson AJ (2012) Aging differentially affects the re-entrainment response of central and peripheral circadian oscillators. J Neurosci 32:16193–16202.

Shan Y, Abel JH, Li Y, Izumo M, Cox KH, Jeong B, Yoo SH, Olson DP, Doyle FJ, and Takahashi JS (2020) Dual-color single-cell imaging of the Suprachiasmatic Nucleus reveals a circadian role in network synchrony. Neuron 108:164–179.e7

Shao X; Somlo S; Igarashi P (2002) Epithelial-specific Cre/lox recombination in the developing kidney and genitourinary tract. J Am Soc Nephrol 13: 1837–1846.

Sinturel F, Gos P, Petrenko V, Hagedorn C, Kreppel F, Storch KF, Knutti D, Liani A, Weitz C, Emmenegger Y, Franken P, Bonacina L, Dibner C, and Schibler U (2021) Circadian hepatocyte clocks keep synchrony in the absence of a master pacemaker in the suprachiasmatic nucleus or other extrahepatic clocks. Genes Dev 35:329–334.

Smyllie NJ, Pilorz V, Boyd J, Meng QJ, Saer B, Chesham JE, Maywood ES, Krogager TP, Spiller DG, Boot-Handford R, White MR, Hastings MH, Loudon AS (2016) Visualizing and quantifying intracellular behavior and abundance of the core circadian clock protein PERIOD2. Curr Biol 26:1880–1886.

Stanewsky R, Kaneko M, Emery P, Beretta B, Wager-Smith K, Kay SA, Rosbash M, and Hall JC (1998) The *cry^b^* mutation identifies cryptochrome as a circadian photoreceptor in *Drosophila*. Cell 95:681–692.

Stokkan KA, Yamazaki S, Tei H, Sakaki Y, and Menaker M (2001) Entrainment of the circadian clock in the liver by feeding. Science 291:490–493.

Stratmann M, Suter DM, Molina N, Naef F, and Schibler U (2012) Circadian *Dbp* transcription relies on highly dynamic BMAL1-CLOCK interaction with E boxes and requires the proteasome. Mol Cell 48:277–287.

Tahara Y, Kuroda H, Saito K, Nakajima Y, Kubo Y, Ohnishi N, Seo Y, Otsuka M, Fuse Y, Ohura Y, Komatsu T, Moriya Y, Okada S, Furutani N, Hirao A, Horikawa K, Kudo T, and Shibata S (2012) *In vivo* monitoring of peripheral circadian clocks in the mouse. Curr Biol 22:1029–1034.

Tam SKE, Brown LA, Wilson TS, Tir S, Fisk AS, Pothecary CA, van der Vinne V, Foster RG, Vyazovskiy VV, Bannerman DM, Harrington ME, Peirson SN (2021) Dim light in the evening causes coordinated realignment of circadian rhythms, sleep, and short-term memory. Proc Natl Acad Sci 118:e2101591118. doi: 10.1073/pnas.2101591118.

van der Vinne V, Martin Burgos B, Harrington ME, and Weaver DR (2020) Deconstructing circadian disruption: Assessing the contribution of reduced peripheral oscillator amplitude on obesity and glucose intolerance in mice. J Pineal Res 69:e12654.

van der Vinne V, Swoap SJ, Vajtay TJ, and Weaver DR (2018) Desynchrony between brain and peripheral clocks caused by CK1 / disruption in GABA neurons does not lead to adverse metabolic outcomes. Proc Natl Acad Sci U S A 115:E2437–E2446.

Weaver DR, van der Vinne V, Giannaris EL, Vajtay TJ, Holloway KL, and Anaclet C (2018) Functionally complete excision of conditional alleles in the mouse suprachiasmatic nucleus by *Vgat-IRES-Cre*. J Biol Rhythms 33:179–191.

Weger M, Weger BD, Diotel N, Rastegar S, Hirota T, Kay SA, Strahle U, and Dickmeis T (2013) Real-time *in vivo* monitoring of circadian E-box enhancer activity: a robust and sensitive zebrafish reporter line for developmental, chemical and neural biology of the circadian clock. Dev Biol 380:259–273.

Welsh DK, Yoo SH, Liu AC, Takahashi JS, Kay SA (2004) Bioluminescence imaging of individual fibroblasts reveals persistent, independently phased circadian rhythms of clock gene expression. Curr Biol 14:2289–2295.

West AC and Bechtold DA (2015) The cost of circadian desynchrony: Evidence, insights and open questions. Bioessays 37:777–788.

Xie X, Kukino A, Calcagno HE, Berman AM, Garner JP, and Butler MP (2020) Natural food intake patterns have little synchronizing effect on peripheral circadian clocks. BMC Biol 18:160–7.

Yagita K, Okamura H (2000) Forskolin induces circadian gene expression of rPer1, rPer2 and dbp in mammalian rat-1 fibroblasts. FEBS Lett 465:79–82.

Yamaguchi S, Kobayashi M, Mitsui S, Ishida Y, van der Horst, G. T., Suzuki M, Shibata S, and Okamura H (2001) View of a mouse clock gene ticking. Nature 409:684.

Yamaguchi Y, Suzuki T, Mizoro Y, Kori H, Okada K, Chen Y, Fustin JM, Yamazaki F, Mizuguchi N, Zhang J, Dong X, Tsujimoto G, Okuno Y, Doi M, and Okamura H (2013) Mice genetically deficient in vasopressin V1a and V1b receptors are resistant to jet lag. Science 342:85–90.

Yamaguchi Y, Okada K, Mizuno T, Ota T, Yamada H, Doi M, Kobayashi M, Tei H, Shigeyoshi Y, Okamura H (2016). Real-time recording of circadian *Per1* and *Per2* expression in the suprachiasmatic nucleus of freely moving rats. J Biol Rhythms 31:108–111.

Yamanaka Y, Honma S, and Honma K (2008) Scheduled exposures to a novel environment with a running-wheel differentially accelerate re-entrainment of mice peripheral clocks to new light-dark cycles. Genes Cells 13:497–507.

Yamazaki S and Takahashi JS (2005) Real-time luminescence reporting of circadian gene expression in mammals. Methods Enzymol 393:288–301.

Yamazaki S, Numano R, Abe M, Hida A, Takahashi R, Ueda M, Block GD, Sakaki Y, Menaker M, and Tei H (2000) Resetting central and peripheral circadian oscillators in transgenic rats. Science 288:682–685.

Yan L, Miyake S, Okamura H (2000) Distribution and circadian expression of *dbp* in SCN and extra-SCN areas in the mouse brain. J Neurosci Res 59:291–295.

Yang N, Smyllie NJ, Morris H, Gonçalves CF, Dudek M, Pathiranage DRJ, Chesham JE, Adamson A, Spiller DG, Zindy E, Bagnall J, Humphreys N, Hoyland J, Loudon ASI, Hastings MH, Meng QJ (2020) Quantitative live imaging of Venus::BMAL1 in a mouse model reveals complex dynamics of the master circadian clock regulator. PLoS Genet 16: e1008729. doi: 10.1371/journal.pgen.1008729.

Yoo SH, Ko CH, Lowrey PL, Buhr ED, Song EJ, Chang S, Yoo OJ, Yamazaki S, Lee C, and Takahashi JS (2005) A noncanonical E-box enhancer drives mouse *Period2* circadian oscillations *in vivo*. Proc Natl Acad Sci U S A 102:2608–2613.

Yoo SH, Yamazaki S, Lowrey PL, Shimomura K, Ko CH, Buhr ED, Siepka SM, Hong HK, Oh WJ, Yoo OJ, Menaker M, and Takahashi JS (2004) PERIOD2::LUCIFERASE real-time reporting of circadian dynamics reveals persistent circadian oscillations in mouse peripheral tissues. Proc Natl Acad Sci U S A 101:5339–5346.

Yoo SH, Kojima S, Shimomura K, Koike N, Buhr ED, Furukawa T, Ko CH, Gloston G, Ayoub C, Nohara K, Reyes BA, Tsuchiya Y, Yoo OJ, Yagita K, Lee C, Chen Z, Yamazaki S, Green CB, and Takahashi JS (2017) *Period2* 3′-UTR and microRNA-24 regulate circadian rhythms by repressing PERIOD2 protein accumulation. Proc Natl Acad Sci U S A 114:E8855–E8864.

Zhang EE, Liu AC, Hirota T, miraglia LJ, Welch G, Pongsawakul PY, liu X, Atwood A, Huss JW 3rd, Janes J, Su AI, Hogenesch JB, Kay SA (2009) A genome-wide RNAi screen for modulators of the circadian clock in human cells Cell 139: 199–210.

Zhang R, Lahens NF, Ballance HI, Hughes ME, and Hogenesch JB (2014) A circadian gene expression atlas in mammals: implications for biology and medicine. Proc Natl Acad Sci U S A 111:16219–16224.

