## Supplemental Materials (Fig S1-S7, Table S1). for "Cell-type specific circadian bioluminescence rhythms in *Dbp* reporter mice"

**This PDF file includes:**

Figure S1 through S7
Table S1

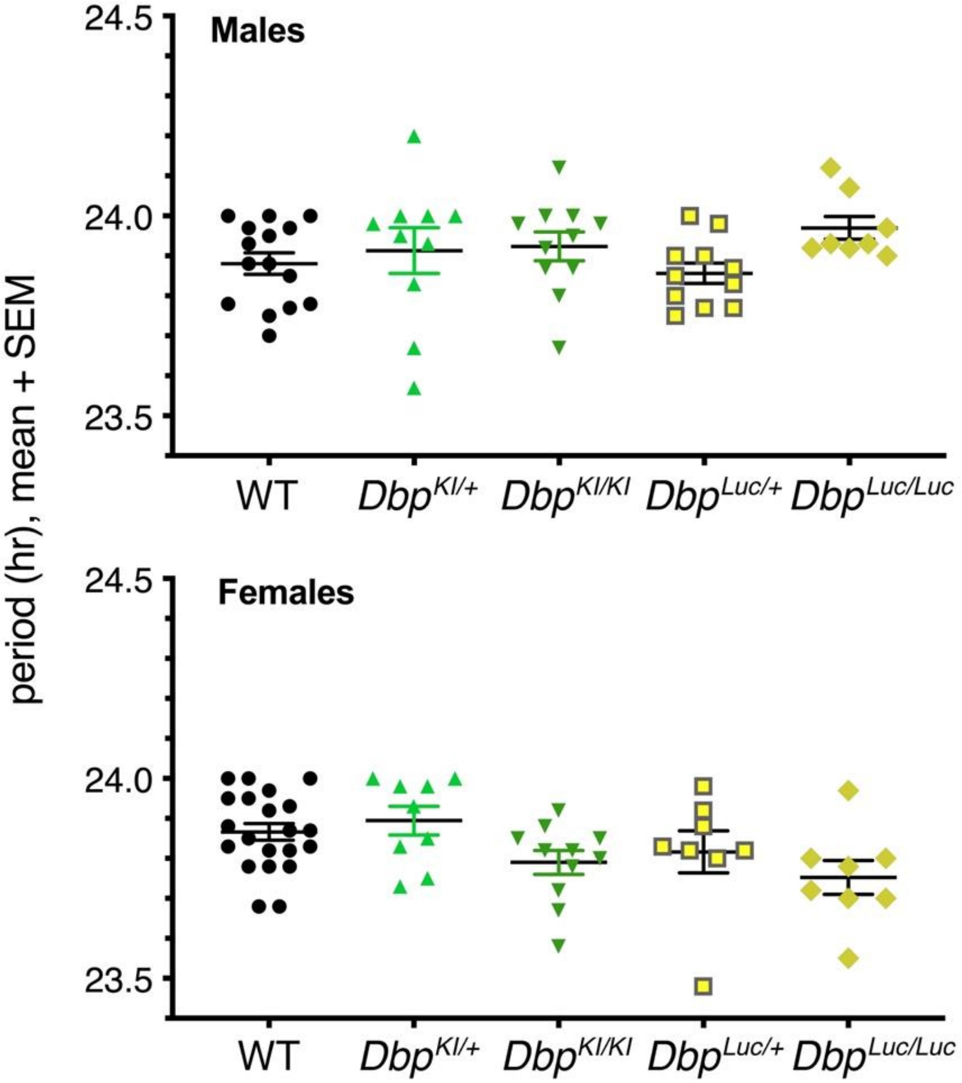


**Fig. S1. Period length values of mice in constant darkness.**

Values are period length values of individual mice, with mean and standard error of the mean also indicted. These data are listed in summary form (mean ± SEM) in Table 1. See text for statistical analysis.

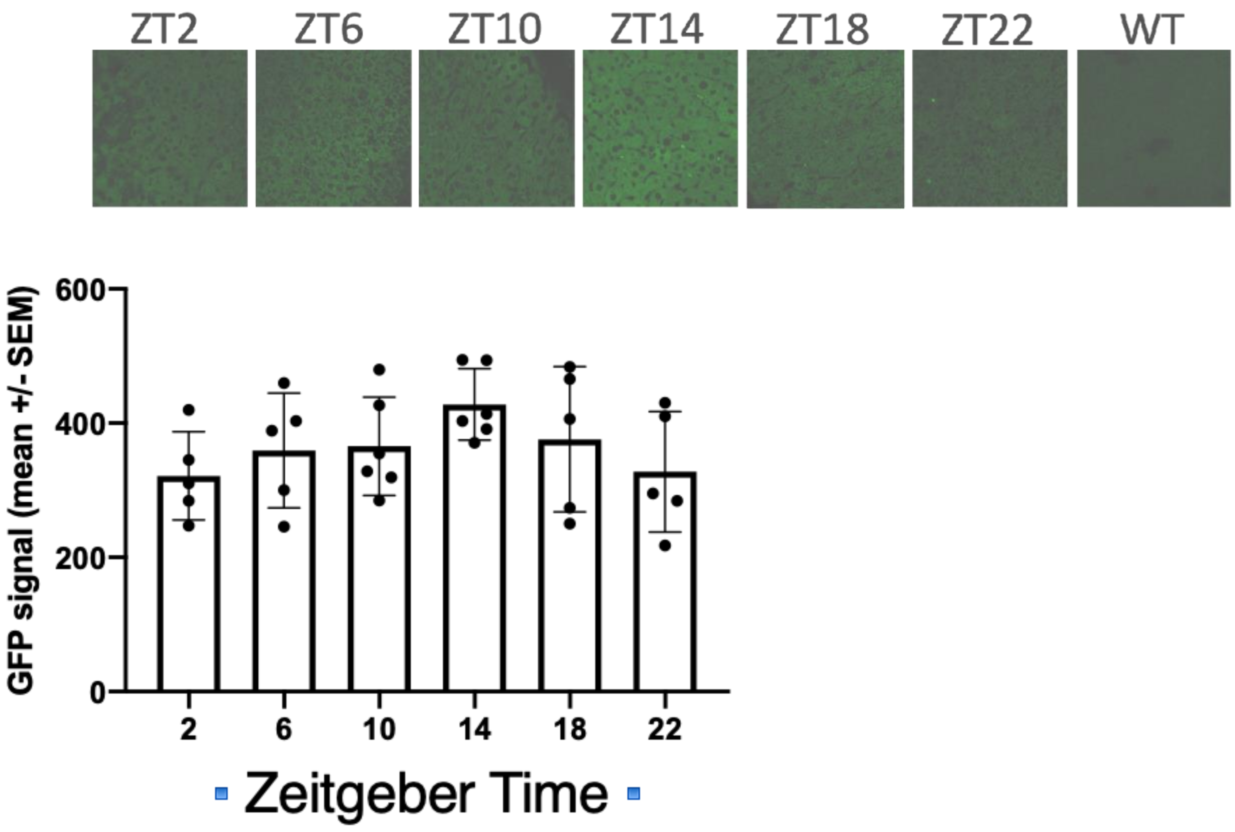


**Figure S2: GFP fluorescence signal in liver sections from *Dbp*$^{KI/+}$ mice.**

Upper panels: Representative images of *Dbp*$^{KI/+}$ liver sections at each timepoint, and in a wild-type (WT) mouse.

Lower panel: Quantification of GFP fluorescence. Each point represents mean fluorescence for the 2-4 sections from 1 mouse.

Methods: Liver tissue was collected from 5-6 *Dbp*$^{KI/+}$ mice (n=5-6 mice per time-point) every 4 hours for 24 hours on the first day in constant darkness. To collect liver tissue, mice were anesthetized with pentobarbital and transcardially perfused with 4% paraformaldehyde and then postfixed for 8 hours. Livers were sectioned at 200µm using a vibratome, and sections mounted on slides with Fluoroshield with DAPI (Sigma-Aldrich). Images were captured at 40x on a Zeiss LSM 880 confocal microscope. ImageJ was used to determine mean fluorescence of 2-4 slices per animal. The average of sections from each of 3 WT mice were 41.2, 64.2 and 66.2 (mean $\pm$ SEM = 57.18 $\pm$ 8.02, n=3 mice).

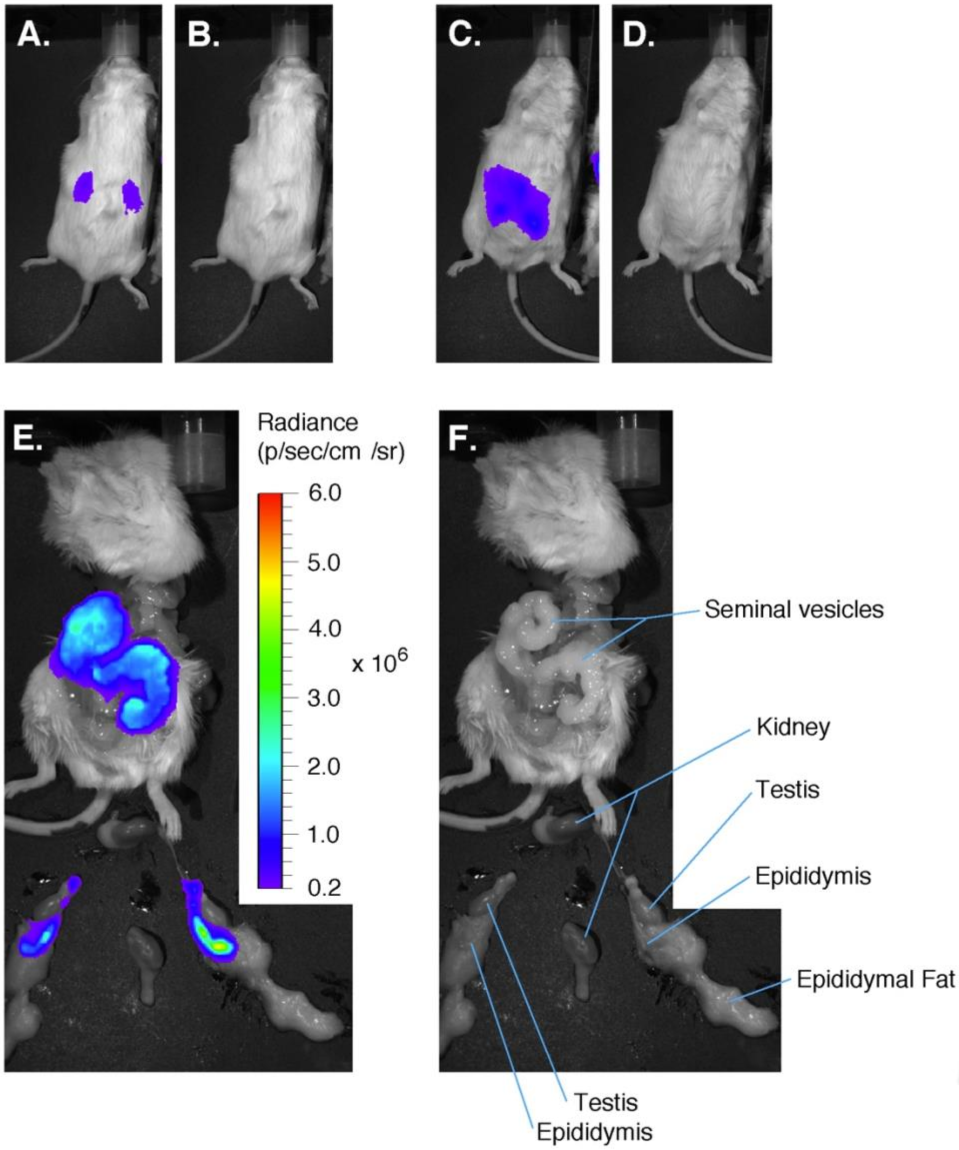


**Figure S3. Bioluminescence distribution in a male 'kidney reporter' mouse.**

Paired bioluminescence images (A,C,E) and photos (B,D,F) from a male albino *Ksp1.3-Cre; Dbp*$^{KI/+}$mouse. The radiance scale inset to Panel E applies to panels A, C, and E.
A,B: Dorsal view captured 11 minutes after injection of D-luciferin (100 ul at 10 mM).
C,D: Ventral view captured 9 minutes after injection of D-luciferin.
E,F: Postmortem dissection identifying the main sources of bioluminescence as seminal vesicles and epididymis. The image in E was captured 14 minutes after D-luciferin injection, within 3 minute of death induced by intentional anesthetic overdose.
Representative of 9 males.

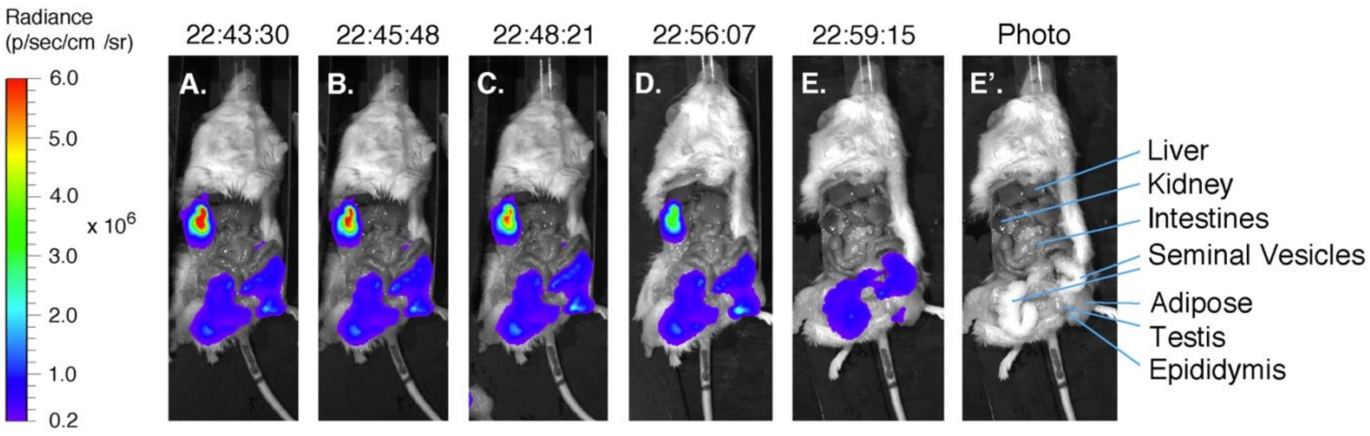


**Figure S4. Postmortem change in bioluminescence distribution in a male kidney reporter.** Luciferin injection (100 ul at 10 mM) at 22:34. The animal was anesthetized but alive in Panels A through D. Panels A-E are 9, 11, 14, 22 and 25 minutes after D-luciferin injection, respectively; clock time is shown above each panel. The radiance scale at left applies to panels A-E. E' is a photo matched to Panel E.

Note the precipitous decline in bioluminescence from the kidney and its preservation from seminal vesicles after the animal died at 22:57 from intentional anesthetic overdose. The animal's left kidney is obscured by liver and other tissues. Taken together with the external images (See Fi.g S3), these results indicate kidney and seminal vesicles are major sources of bioluminescence in male *Ksp1.3-Cre; Dbp*$^{KI/+}$ mice.

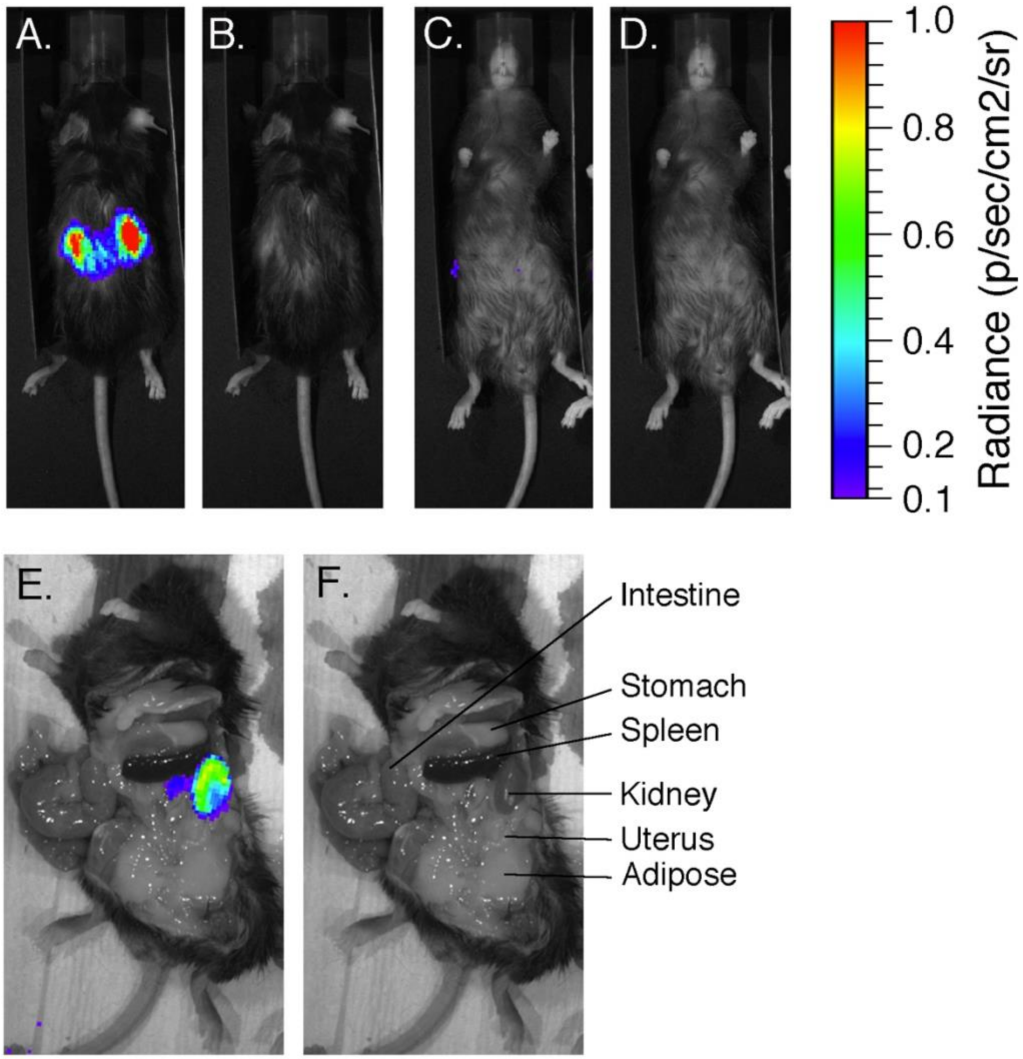


**Figure S5. Biolumnescence originates from the kidney in female ‘kidney reporter’ mice.**

A-D Matched bioluminescence images and photos of intact mice.
A,B: Dorsal view captured 9 minutes after D-luciferin injection (100 ul at 10 mM).
C,D: Ventral view captured 10.5 minutes after D-luciferin injection.
E,F: Postmortem dissection identifies the source of bioluminescence as the kidney. The image was captured 15 minutes after D-luciferin injection, within 3 minute of death induced by intentional anesthetic overdose. The animal’s right kidney is obscured by other abdominal contents. External views representative of 16 females. Dissection representative of 7 females.

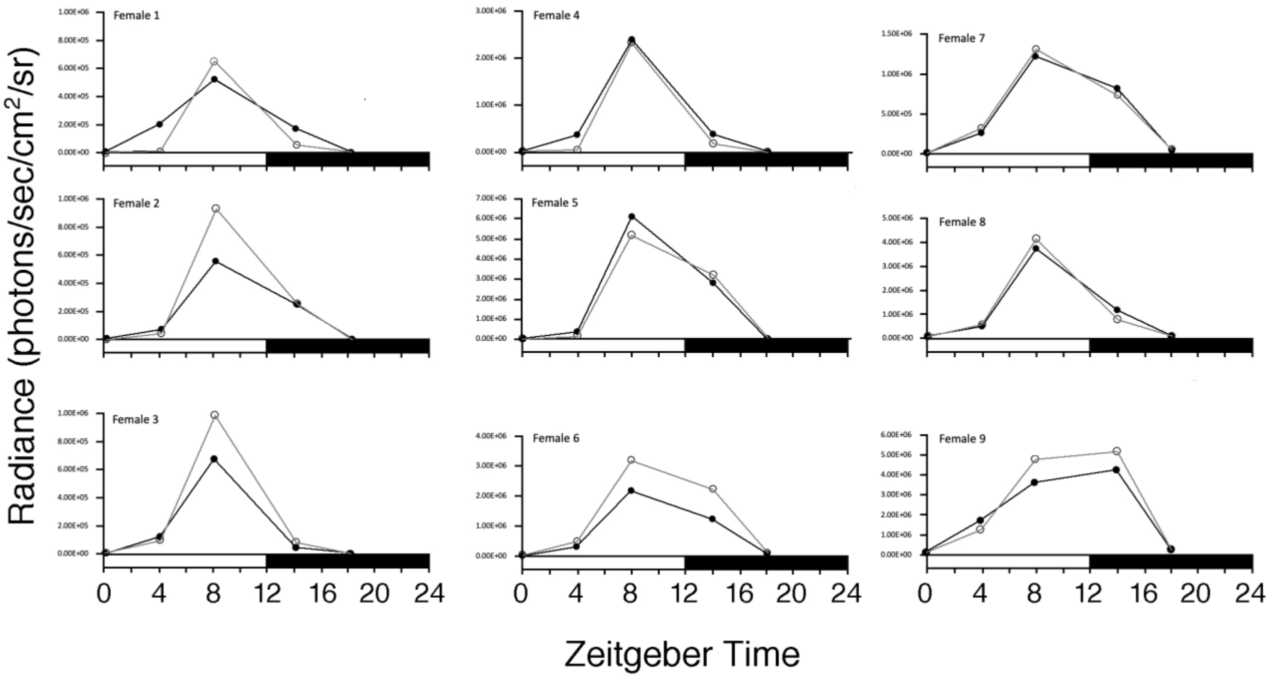


**Figure S6: Bioluminescence rhythms in female kidney reporter mice.**

Quantitative assessment of renal bioluminescene rhythms in intact female kidney reporter mice (albino *Ksp1.3-Cre; Dbp*$^{KI/+}$, n=9). For each female, right kidney (filled symbols, black line) and left kidney(open symbols, gray line) values are shown. Images (dorsal view) were captured 9 to 9.5 min after injection of D-luciferin (100 ul at 10 mM). Time-points are ZT0-1, 4-5, 8-9, 14-15, and 18-19; data are plotted at the beginning of each interval. In 8 of 9 mice, the highest level of bioluminescence occurred at ZT8. Friedman one-way analysis of variance indicated a significant difference among the timepoints ($F_r$ =32.71, k=5, n=9, p<0.0001). Dunn's test revealed that the ranks at the ZT8 timepoint differed significantly from the ranks at ZT0 and ZT18 (multiplicity-corrected, two-tailed p-value < 0.0005), with a trend toward a difference at ZT4 (multiplicity-corrected, two-tailed p-value = 0.0684). Ranks at ZT8 and ZT14 did not differ significantly.
Methods: Radiance was assessed using Living Image software for ROI's of fixed size for each kidney at the five Zeitgeber time-points. For each kidney, the rank order of bioluminescence levels among the five timepoints was determined. When the order was not identical for both kidneys within an animal (as occurred twice), the bioluminescence values from the two kidneys at each time-point were averaged; the ranking of timepoints was then determined using the average values. Thus, bioluminescence from both kidneys was considered, but each animal contributed only one set of rankings for statistical analysis.

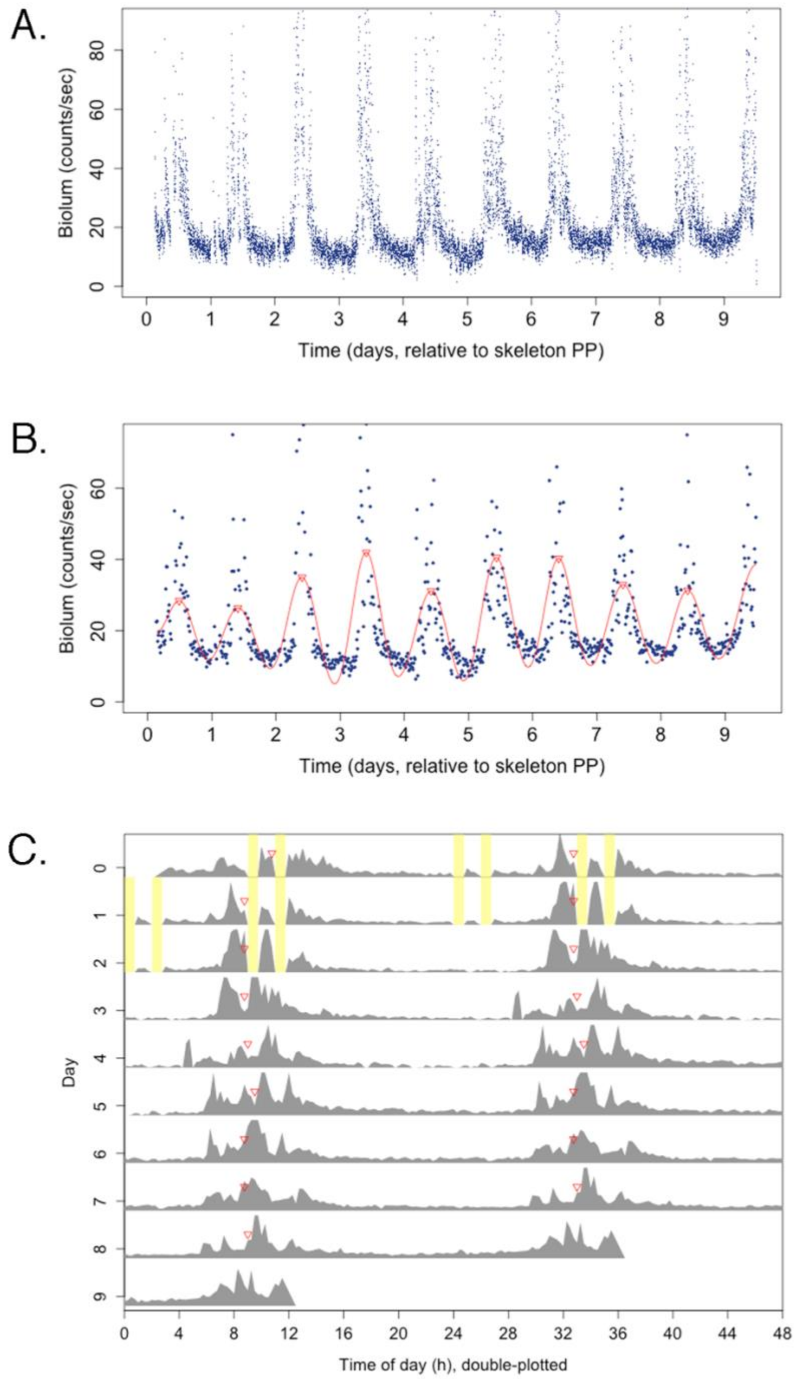


**Figure S7. Bioluminescence Rhythms from a Liver Reporter Mouse.** Panels A,B and C represent the same data, from a female mouse with 2 mM D-luciferin in its drinking water. Panel A shows raw bioluminescence data, excluding outliers above the 98th percentile. Panel B shows 15-minute median binned values as blue dots, the DWT circadian component plus smooth as a red curve, and the DWT-calculated time of peak as red triangles. Panel C displays the bioluminescence data (gray) in actogram format, with the skeleton photoperiod in yellow and red triangles marking time of peak for each day. Time zero is set to the start of the first dawn light pulse of the skeleton photoperiod (skeleton PP) during bioluminescence recording.

**Table S1. PCR Primer Pairs.**

**1. Primer Pairs for Confirming the Targeted *Dbp* Locus**

Primer pair C (Internal to the construct; forward in GFP, reverse in Luc2)

| | |
|---|---|
| *Dbp*-C- Forward | 5′- CAAGGTGAACTTCAAGATCCGC -3′ |
| *Dbp*-C-Reverse | 5′- CCCATGCTGTTCAGCAGCTCG -3′ |

Primer pair F  (spans 5′ end;  forward in intron 3 outside construct, reverse in T2A sequence)

| | |
|---|---|
| *Dbp*-F- Forward | 5′ - GGAAGCATCTTTTCCAGCTGG - 3′ |
| *Dbp*-F-Reverse | 5′ - TTCCTCTGCCCTCTCCACTGC - 3′ |

Primer Pair H (spans 3′ end; forward in Luciferase, reverse in 3′ UTR outside construct)

| | |
|---|---|
| *Dbp*-H- Forward | 5′ - GATTCTCATTAAGGCCAAGAAGG - 3′ |
| *Dbp*-H- Reverse | 5′ - CATGGCGAGTTGGTGGAACCAGC - 3′ |

Primer Pair ‘confirm’ (external to and spanning the entire construct)
(forward in intron 3 outside the construct, reverse in 3′ UTR outside the construct)

| | |
|---|---|
| *Dbp*-span-Forward | 5′ - GATGTGTCCTAACAAGCTGGAGC - 3′ |
| *Dbp*-span-Reverse | 5′ - AAGCCACAAGCCTGAACGAGC - 3′ |

**2. Genotyping Primer sets**

*Dbp* Primer set “4A” (Common forward primer in exon 4, allele-specific reverse primers)

| | |
|---|---|
| *Dbp*-4A-Forward | 5′ - TGCTGTGCTTTCACGCTACCAGG - 3′ |
| *Dbp*-4A-Reverse in GFP | 5′ - AGTCGTGCTGCTTCATGTGGTCG - 3′ |
| *Dbp*-4A-Reverse in Luc2 | 5′ - TCGTTGTAGATGTCGTTAGCTGG - 3′ |
| *Dbp*-4A-Reverse in 3’UTR | 5′ - TTCAGGATTGTGTTGATGGAGGC - 3′ |

Primer set *Clock/Cre* {Clock J (internal control) plus Cre-370, used as a 4-primer mix}

| | |
|---|---|
| *Clock* Forward | 5′- GCAAGAAGAACTAAGGAAAATTCAA-GAGCAACTTCAGATGGTCCATGGTCAA-GGGCTACAGTT - 3′ |
| *Clock* Reverse | 5′- TAGTGCCCTAGATGGCCCTGTTGG -3′ |
| Cre-370 Forward | 5′ - ACCTGAAGATGTTCGCGATTATCT - 3′ |
| Cre-370 Reverse | 5′ - ACCGTCAGTACGTGAGATATCTT - 3′ |

Per2::LUCIFERASE(SV) (Common forward primer, allele-specific reverse primers)

| | |
|---|---|
| *Per2* Common Forward | 5′ - CTGCGAGAGTGAGGAGAAAGGC - 3′ |
| *Per2* WT-specific Reverse | 5′ - GGATTTCCTCCTAAACCTCCC - 3′ |
| *Per2* Luc-specific Reverse | 5′ - GTAGATGAGATGTGACGAACG - 3′ |

**3. Primer pairs for Generating DIG-labeled Probes**

| | |
|---|---|
| *Actin* Forward: | 5′ - TCAGAAGGACTCCTATGTGGG - 3′ |
| *Actin* Reverse: | 5′ - GATCCACACAGAGTACTTGCG - 3′ |
| *Dbp* Forward: | 5′ - AATGACCTTTGAACCTGATCCC - 3′ |
| *Dbp* Reverse: | 5′ - TCACAGTGTCCCATGCTGGG - 3′ |
| *GFP* Forward: | 5′ - CTGAAGTTCATCTGCACCACCG - 3′ |
| *GFP* Reverse: | 5′ - GTGCTCAGGTAGTGGTTGTCGG - 3′ |
| *Luc2* Forward: | 5′ - GCTTCGAGGAGGAGCTATTCTTGC - 3′ |
| *Luc2* Reverse: | 5′ - CAGCAGGATGCTCTCCAGTTCGG - 3′ |
